# RNAi epimutations conferring antifungal drug resistance are inheritable

**DOI:** 10.1101/2024.10.15.618540

**Authors:** Carlos Pérez-Arques, María Isabel Navarro-Mendoza, Ziyan Xu, Grit Walther, Joseph Heitman

## Abstract

Epimutations modify gene expression and lead to phenotypic variation while the encoding DNA sequence remains unchanged. Epimutations mediated by RNA interference (RNAi) and/or chromatin modifications can confer antifungal drug resistance and may impact virulence traits in fungi. However, whether these epigenetic modifications can be transmitted across generations following sexual reproduction was unclear. This study demonstrates that RNAi epimutations conferring antifungal drug resistance are transgenerationally inherited in the human fungal pathogen *Mucor circinelloides*. Our research revealed that RNAi-based antifungal resistance follows a DNA sequence-independent, non-Mendelian inheritance pattern. Small RNAs (sRNAs) are the exclusive determinants of inheritance, transmitting drug resistance independently of other known repressive epigenetic modifications. Unique sRNA signature patterns can be traced through inheritance from parent to progeny, further supporting RNA as an alternative molecule for transmitting information across generations. Understanding how epimutations occur, propagate, and confer resistance may enable their detection in other eukaryotic pathogens, provide solutions for challenges posed by rising antimicrobial drug resistance (AMR), and advance research on phenotypic adaptability and its evolutionary implications.

## Introduction

The widespread emergence of antimicrobial drug resistance (AMR) is threatening global health, undermining the past century’s advances that revolutionized modern medicine and transformed the treatment of infectious diseases (1). Understanding the molecular mechanisms that confer and transmit resistance is essential, not only for developing novel antimicrobial therapies but also for diagnosing and countering the rise of resistant pathogens. Traditionally, AMR studies have focused on bacteria and antibiotics, yet the threat extends to eukaryotic microbes as well. Fungal infections are increasing at alarming rates, and therapeutic options are scarce. This has prompted the World Health Organization (WHO) to prioritize research and development into the mechanisms of multi- and pan-resistance in fungal pathogens (2), which has recently been termed fAMR (3).

Antifungal drug resistance typically arises from genetic mutations that compromise the interaction of the drug with its target (4). These changes in the DNA sequence are stable and follow the laws of Mendelian inheritance. Aneuploidy, chromosomal rearrangements, and copy number variation can also mediate drug resistance, tolerance, heteroresistance, and persistence (5). However, these genomic changes are unstable and transient, usually reverting once drug selective pressure ceases. But there are other, seemingly silent mechanisms that operate —ones that alter gene expression without changing the DNA sequence. These are called epimutations, and rely upon epigenetic mechanisms such as histone modification, DNA methylation, or RNA interference (RNAi) to cause phenotypic changes (6–9). Fungal epimutations based on RNAi were first identified in *Mucor* species, showing that small RNAs (sRNAs) silencing of the FK506 drug target resulted in epigenetic, reversible resistance (6). Similarly in fission yeast, heterochromatin silencing mediated by histone H3 lysine 9 methylation (H3K9me) can result in caffeine resistance (8). These epimutations cause transient drug resistance that reverts to susceptibility after several mitotic divisions without the drug. However, whether epimutations can be transmitted across generations —a process known as transgenerational inheritance— and the mechanisms driving their heritability, remain subjects of ongoing debate (10–12).

In this study, we show how spontaneous RNAi epimutations conferring antifungal drug resistance are transmitted to the next generation after sexual reproduction in the human fungal pathogen *Mucor circinelloides*. The inheritance pattern is DNA sequence-independent and non-Mendelian, relying exclusively on sRNA molecules as the determinants of inheritance, uncoupled from other repressive epigenetic modifications.

## Results

### RNAi epimutations in opposite mating types of *M. circinelloides*

RNAi epimutations conferring fAMR were first reported in *Mucor circinelloides* formae *lusitanicus* and *circinelloides* (6), recently identified as separate phylogenetic species (PS10 and PS14) (13). However, neither species can produce viable progeny through sexual reproduction, presenting a challenge to study genetic and epigenetic inheritance. Therefore, we studied epimutations in two fertile, opposite mating types (– and +) of *M. circinelloides* <u>p</u>hylogenetic species <u>15</u> (referred to as PS15). PS15 was selected as it is the only species among 16 in the *Mucor circinelloides* complex able to complete the sexual cycle under laboratory conditions (13). Despite this advantage, epimutation, RNAi proficiency, and meiotic recombination had not been assessed in this species, and it was unclear if the observed progeny resulted from bona fide sexual reproduction. To address these limitations, whole-genome assemblies and gene annotations for both opposite mating types (PS15– and PS15+) were generated with Oxford Nanopore Technologies (ONT) long reads (Extended Data Table 1). Each genome assembly consists of ∼ 37 Mb distributed among fourteen and fifteen contigs, showing an N50 of 3.29 and 2.90 Mb for PS15– and PS15+, respectively. Both assemblies exhibit a high level of completeness, with 97.3% of conserved eukaryotic genes present. Protein-protein similarity searches identified homologs of the principal RNAi components involved in establishing and maintaining epimutations (6, 14–16) (Extended Data Fig. 1a). sRNA sequencing of both PS15– and + confirmed the presence of canonical small interfering RNAs (siRNAs), characterized by discrete lengths of 21 to 24 nt and predominantly a 5’ uracil (16) (Extended Data Fig. 1b).

After confirming RNAi proficiency, RNAi-targeted *fkbA* epimutants were isolated by challenging PS15– and + fungal spores with FK506, both alone and in combination with rapamycin (Fig. 1a and Extended Data Fig. 1c, d). Both drugs target FKBP12 encoded by the *fkbA* gene (17), causing similar fungistatic effects: FK506 enforces yeast-like, decreased growth by inhibiting the calcineurin pathway, whereas rapamycin compromises growth by inhibiting TOR signaling (Extended Data Fig. 1c). This dual drug approach enabled efficient screening for loss of FKBP12 function. By this approach, 14 FK506-resistant isolates were obtained after five days of continuous exposure to the drug (Extended Data Fig. 1d). Seven PS15– (E1– to E6– and M1–) and six PS15+ isolates (E10+, E12+, E15+, M1+ to M3+) were selected for their resistance to FK506 and rapamycin (Fig. 1b, Extended Data Fig. 1e, and Extended Data Table 2), indicating loss of FKBP12 function. Sanger-sequencing revealed *fkbA* missense mutations in four isolates, M1– and M1+ to M3+ (and so named <u>M</u>utants, Extended Data Fig. 1f). No DNA sequence changes were detected within the *fkbA* coding or untranslated regions of the remaining nine isolates, E1– to E6–, or E10+, E12+, and E15+ (Extended Data Fig. 1f). Follow-ing mitotic passage without drug, these resistant isolates reverted to wildtype susceptibility, indicating resistance is unstable and transient (Fig. 1c). Because a similar unstable resistance due to RNAi was previously reported in other *Mucor* species (6, 18), northern blot analyses were conducted, revealing antisense sRNAs targeting *fkbA* in these unstable resistant colonies during active epimutation, i.e. before their reversion. These sRNAs were undetectable in wildtype and *fkbA* mutant negative control strains, suggesting epimutation-driven resistance (<u>E</u>pimutants E1– to E6–, E10+, E12+, and E15+, Extended Data Fig. 1g).

**Fig. 1.**
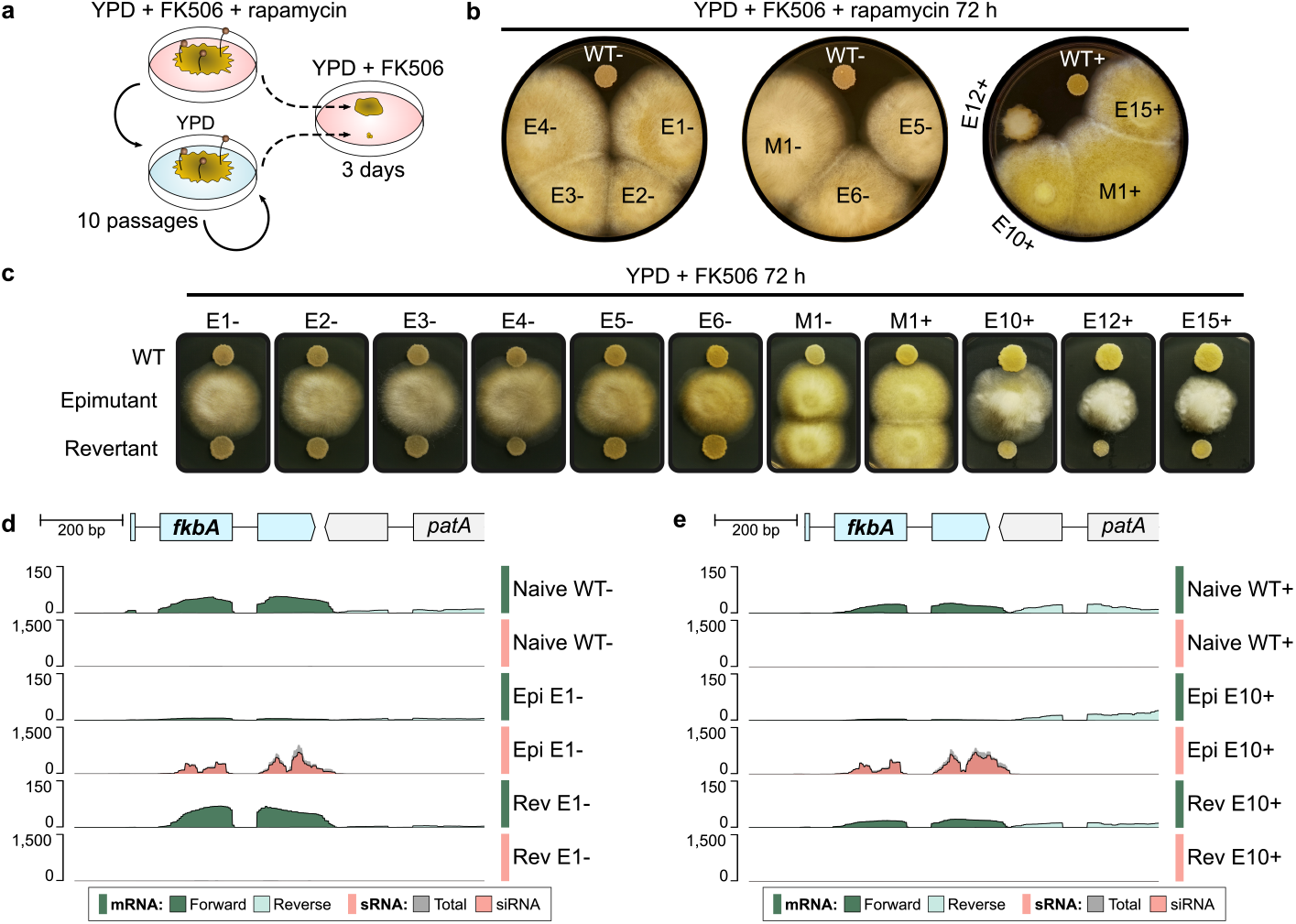
Two opposite mating types from *Mucor circinelloides* <u>P</u>hylogenetic <u>S</u>pecies <u>15</u> (PS15) exhibit RNAi-based epigenetic resistance to FK506 and rapamycin. **(a)** Experimental design to validate FK506 reversible resistance due to loss of FKBP12 function. Briefly, epimutations were induced by exposing wildtype, naïve spores to FK506 for five days, as shown in Extended Data Fig. 1d. Resistant mycelium colonies were transferred to FK506 and rapamycin medium to confirm loss of FKBP12 function. Confirmed resistant isolates were serially passaged in drug-free medium for ten cycles of vegetative growth. Spores collected from the final passage were rechallenged with FK506 to assess resistance stability compared to the original resistant isolate. **(b)** Isolates from PS15 – and + mating types exhibiting FK506 resistance were challenged with both FK506 and rapamycin for 72 hours, comparing growth to wildtype strains (WT). Isolates resistant to both drugs are labelled in black, and susceptible isolates in white. Resistant isolates with wildtype *fkbA* DNA sequence are labeled as <u>E</u>pimutants (E); those harboring *fkbA* mutations are labeled as Mutants (M). **(c)** Epimutants and mutants were passaged on non-selective medium for ten 84-hour passages (revertants) and transferred onto FK506-selective medium with wildtype sensitive and resistant control isolates to assess reversion. **(d, e)** Genomic plots of the *fkbA* locus (1 kb) depicting the *fkbA* gene (light blue) and 3’ region of the neighboring *patA* gene (light gray). Stranded messenger RNA (mRNA) and sRNA coverage tracks are color-coded for a representative <u>E</u>pimutant, and its corresponding <u>WT</u> control and <u>Rev</u>ertant isolate from the **(d)** – and **(e)** + mating type. Stranded mRNA coverage was aligned to the forward and reverse strands and overlaid in different shades of green. sRNA coverage shows all aligning reads (total, dark gray) overlaid with reads exhibiting typical small interfering RNA features (siRNA, red), i.e., antisense 21-24 nt reads with a 5’-uracil.

RNAi epimutations were further analyzed by sRNA and ribosomal RNA (rRNA)-depleted RNA sequencing. sRNA-producing loci were identified in PS15 naïve wildtype strains, revealing that sRNAs derive primarily from genes and repeated sequences (Extended Data Fig. 2a and Supplementary Data 1), and the most active siRNA loci arise from clusters of repeated elements (Extended Data Fig. 2b). Low levels of sRNAs were detected across the *fkbA* locus in wildtype strains (Extended Data Fig. 2b, c). However, these sRNAs lack canonical siRNA features, such as enrichment in lengths from 21-24 nt length enrichment or 5’-uracil bias, and are predominantly sense to the *fkbA* mRNA (Extended Data Fig. 2d, e), compatible with transcript degradation (16). In contast, the resistant isolates showed an accumulation of *fkbA* antisense siRNAs, accompanied by a marked decrease in *fkbA* mRNA levels, consistent with posttranscriptional gene silencing. Upon removal of drug selective pressure, revertant isolates lost siRNAs and *fkbA* mRNA was restored to wildtype levels (Fig. 1d, e). Our results demonstrate that RNAi-epimutations conferring FK506 and rapamycin resistance are not limited to *M. lusitanicus* and *M. circinelloides* PS14, but are also reproducibly induced in *M. circinelloides* PS15, highlighting the widespread relevance of this adaptive mechanism across the *Mucor circinelloides* species complex.

### Non-Mendelian inheritance of epigenetic drug resistance after sexual reproduction

Previous studies (6, 18) and this work shows that RNAi epimutations in *Mucor* species are stable enough to be transmitted through multiple rounds of mitotic division before reversion. Therefore, we hypothesized that epimutations might be similarly transmitted through meiosis and sexual reproduction without drug selective pressure. To test epimutation heritability and determine the genetic basis of the resistance trait, we obtained F1 progeny from a series of genetic crosses. First, an epimutant and a wild type were crossed, utilizing epimutants from both mating types (Fig. 2a, b). Second, two epimutants from opposite mating types were crossed (Fig. 2c). After successful mating of haploid, opposite-mating type mycelia, the reproductive structures known as zygospores were formed (19). Zygospores from each cross were individually dissected and germinated after a dormancy period of 2 to 8 weeks, exhibiting germination rates from 1 to 2%. Within each zygospore, haploid nuclei fuse into a diploid nucleus that undergoes meiosis (20, 21). Only one of the four meiotic products is mitotically amplified to form a germsporangium containing genetically identical germspores (22, 23), which were considered F1 progeny.

**Fig. 2.**
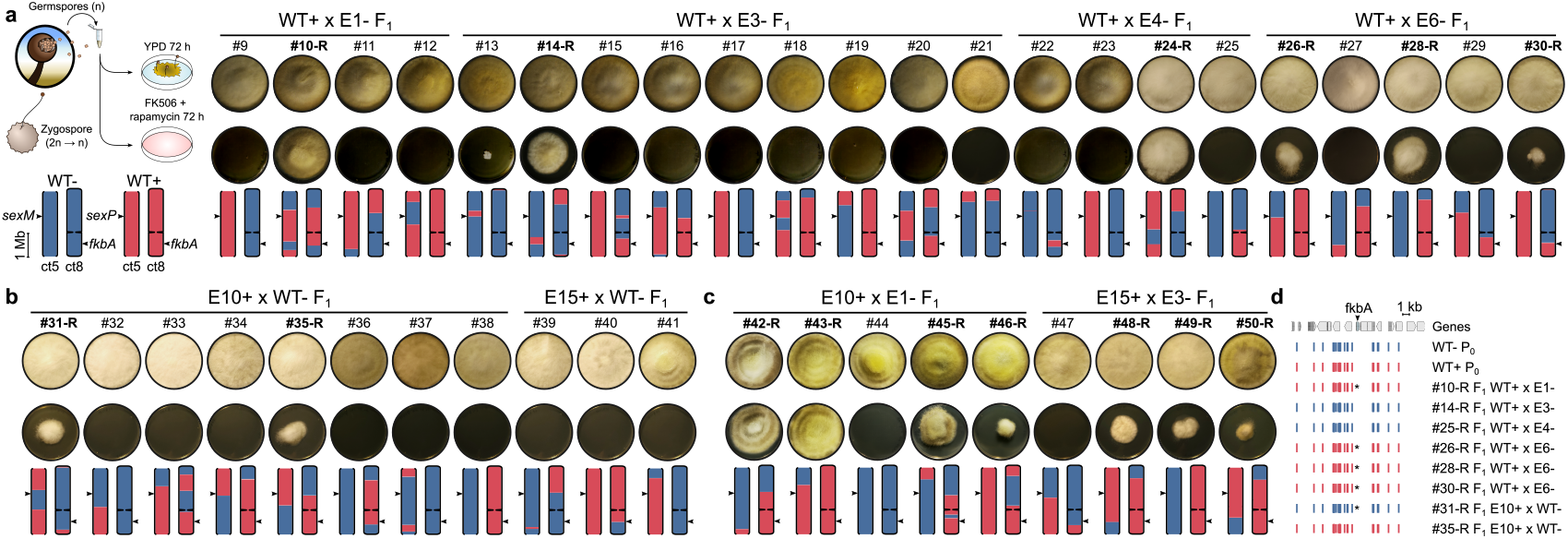
FK506 resistance displays non-mendelian inheritance ratios in progeny from sexual reproduction. **(a-c)** *M. circinelloides* PS15 progeny from sexual crosses are shown: **(a)** – mating type epimutant crossed with + mating type wildtype strain, **(b)** + mating type epimutant crossed with – mating type wildtype strain, and **(c)** + mating type epimutant crossed with – mating type epimutant. Growth on YPD as a viability control and on FK506+Rapamycin medium is shown for each progeny, along with two contigs containing the *sex* (ct5) and *fkbA* (ct8) loci, indicated by arrows. The contig drawings depict the inheritance pattern of SNVs (– mating type in blue and + mating type in red), demonstrating meiotic recombination and/or independent chromosome assortment. Ct5 is depicted as an incomplete chromosome arm lacking telomeric repeats (open ends), whereas ct8 is a full chromosome depicting a centromere (constriction) and telomeric repeats on both ends (closed ends). **(d)** Genomic plot showing a 20 kb region centered on the *fkbA* gene. Genetic variation (shown as SNVs) inherited from either mating-type wildtype parent (WT P_0_) is shown for each resistant (−R) progeny isolated from the crosses of epimutants with wildtype.

Drug susceptibility to FK506 alone (Extended Data Fig. 3a-c) and in combination with rapamycin (Fig. 2a-c) was assessed in the F1 progeny without any intermediate passaging. Resistance was considered inherited if observed in both conditions. This implies the resistance was present in the progeny prior to the drug challenges, rather than arising from two independent de novo epimutations due to drug exposure. The cross with the – mating type epimutant yielded 6 out of 22 resistant progeny (Fig. 2a), which is inconsistent with a 1:1 inheritance ratio expected from Mendelian inheritance of a monogenic trait in haploid organisms [goodness-of-fit (GOF) Chi-square test, *p*-val 0.03301]. Similarly, the + mating type epimutant yielded a 2 out of 11 resistance inheritance ratio (Fig. 2b, GOF Chi-square test, *p*-val 0.03481). Together, these results indicate that resistance can be inherited from either mating type at similar rates, ruling out mating type bias or uniparental inheritance. Importantly, resistance was inherited at significantly lower ratios than those expected from Mendelian inheritance, supporting RNAi epimutations as epigenetic traits. The remaining progeny showed wildtype susceptibility to FK506, suggesting the epimutation was either lost or not inherited. On the other hand, crossing two epimutants yielded a higher frequency of drug-resistant progeny (7 out of 9, Fig. 2c) but still inconsistent with a 100% inheritance expected from a Mendelian monogenic trait present in both haploid parents (GOF Chi-square test, *p*-val = 0.01776). Control crosses between two naïve wild-type isolates yielded only susceptible progeny (8 out of 8, Extended Data Fig. 3d). Similarly, F1 progeny from an *fkbA* mutant showed resistance ratios consistent with Mendelian inheritance (3 out of 5, Extended Data Fig. 3e), demonstrating that mutations and epimutations affecting *fkbA* show distinct inheritance rates.

Molecular analysis demonstrated that the progeny isolated from these crosses resulted from bona fide sexual reproduction including meiosis. We analyzed genetic variation across the PS15– and PS15+ genomes, identifying a total of 18,731 single nucleotide variants (SNVs) across 13 genomic blocks that exhibited full synteny and contiguity in both opposite mating types (Extended Data Fig. 4). These synteny blocks include six telomere-to-telomere, complete chromosomes and seven contigs, with nine annotated centromeres. By identifying inherited genetic variation across these synteny blocks, we were able to show meiotic recombination events and independent chromosome assortment in all of the F1 progeny isolates (Fig. 2a-c and Extended Data Fig. 5).

The distribution of the inherited SNVs revealed the source of the *fkbA* allele in the progeny. Resistant progeny inherited the *fkbA* locus at ratios consistent with Mendel’s laws (5:3, Chi-square test, *p*-val 0.4795). The results demonstrate that the epimutation resistance is not genetically linked to the *fkbA* locus, as half of the progeny that inherited resistance from the epimutant parent inherited the *fkbA* allele from the naïve wildtype parent (Fig. 2d, asterisks). Similarly, susceptible progeny inherited the *fkbA* allele from either parent, regardless of their susceptibility. Whole-genome analysis of parents and progeny revealed no changes in the *fkbA* DNA sequence or chromosomal rearrangements contributing to the observed resistance (Extended Data Fig. 6), supporting epigenetic modifications as the exclusive cause of epimutational resistance.

Similarly, in-depth variant calling was conducted to identify off-target mutations that might contribute to FK506 and rapamycin resistance in the epimutant parental strains (E1– to E6–, E10+, E12+, and E15+). Variants were manually curated to identify false positives, discarding variants present in the wildtype negative controls (Extended Data Fig. 7 and Supplementary Data 2). As expected, mutations in *fkbA* were identified in the M1-(TC>T) and M1+ (AGG>A) positive control strains, confirming previous Sanger sequencing results and validating the pipeline. Notably, the epimutant E12+ harbored a substitution (T>C) in a non-coding region upstream of PS15p_212321, which encodes a predicted DNA transposon and is therefore unlikely to be relevant to FK506 and rapamycin epimutational resistance.

### Epimutation inheritance depends on sRNAs, uncoupled from heterochromatin formation

Substantial evidence supports epigenetic inheritance across a range of eukaryotes, including animals, plants, fungi (12), and other eukaryotic microorganisms (24). This inheritance mainly relies on chromatin modifications, such as H3K9me, H3K27me, and/or DNA 5-methylcytosine (5mC). Although *Mucor* spp. lack the necessary components for H3K27me and 5mC (25– 27), heterochromatin can form and be maintained through H3K9me (26). Therefore, we hypothesized that RNAi and H3K9me could be coupled to stabilize epimutations, allowing their inheritance in the progeny. This hypothesis was tested by chromatin immunoprecipitation (ChIP) targeting H3K9me2 in PS15 epimutant parents and progeny.

H3K9me2-based heterochromatin is abundant in repeats, particularly at centromeres and telomeres (Fig. 3a), consistent with previous findings in *Mucor lusitanicus* PS10 (26). No H3K9me2 was detected at either the *fkbA* locus in epimutant parents of either mating type or their drug-resistant progeny (Fig. 3a, b and Extended Data Fig. 8a). Instead, abundant siRNAs target *fkbA* in these epimutant isolates, and this results in a decrease in *fkbA* mRNA levels (Fig. 3b). Despite this reduction, *fkbA* transcription remains active during epimutation as mRNA is still detectable (Fig. 3b and Extended Data Fig. 8b), and RNA polymerase II (RNAP) binding across this locus does not differ significantly between naïve and epimutant strains (Fig. 3b-d, Extended Data Fig. 8a, and Supplementary Data 3), indicating that *fkbA* is not transcriptionally silenced during epimutation.

**Fig. 3.**
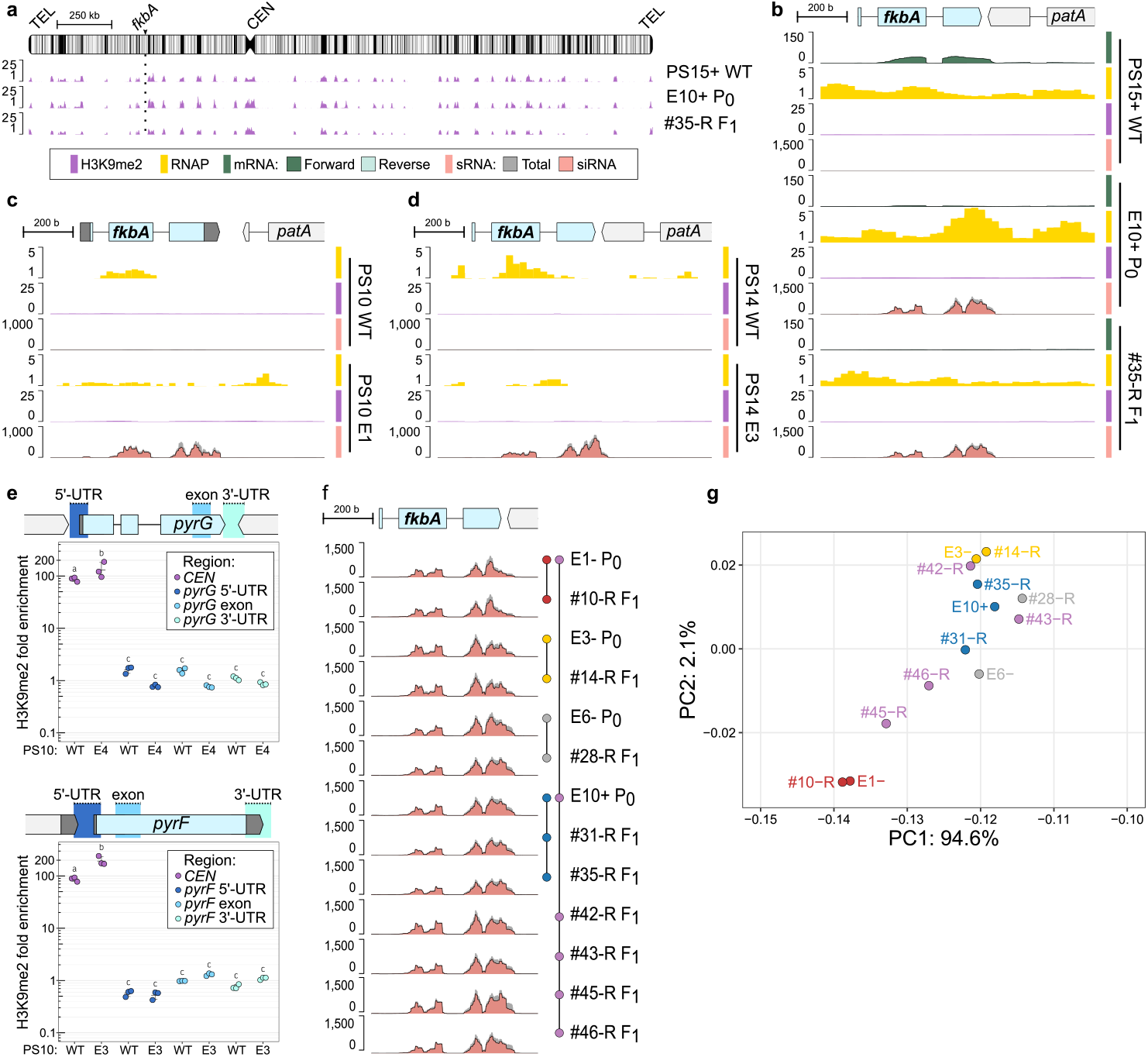
Inherited epigenetic resistance is exclusively driven by RNAi and independent of heterochromatin marks. **(a-d)** Genomic plots showing H3K9me2 enrichment (purple), RNA Polymerase II enrichment (yellow), stranded mRNA (overlaid shades of green for forward and reverse strand), and/or sRNA (dark gray for total sRNA overlaid with red for siRNA) in **(a-b)** *M. circinelloides* PS15 wild type, an epimutant parent (P_0_) and its F_1_ progeny across **(a)** the full chromosome containing the *fkbA* gene (arrowhead and dotted line). Repeats are depicted as black vertical lines, the centromere as a constriction (*CEN*), and telomeric sequences as rounded ends (*TEL*); **(b)** a magnified view of the *fkbA* locus (1 kb); **(c, d)** a wild type (WT) and *fkbA* epimutant (E) from **(c)** *Mucor lusitanicus* PS10 and **(d)** *M. circinelloides* PS14. **(e)** Lack of H3K9me2 ChIP enrichment across *pyrG* and *pyrF* regions is shown for *fkbA* susceptible (WT) and epimutant (EM) *M. lusitanicus* PS10 isolates. For each gene, amplicons corresponding to the 5’-and 3’-UTRs and exonic regions were tested by qPCR, as indicated in the gene model. A highly enriched centromeric amplicon (*CEN*) was also included for comparison. Average values and standard deviations were derived from technical triplicates. Significant differences were determined using a one-way ANOVA and Tukey HSD tests. Average values not showing significant differences are indicated by the same letter. **(f)** The *fkbA* genomic plot shows total and siRNA coverage for a representative set of epimutant parental (P_0_) and F_1_ resistant progeny (#-R) isolates. Parents and progeny are represented by colored circles connected by lines. **(g)** Principal component analysis of the relative abundance of sRNA sequences complementary to the *fkbA* locus in epimutant parents and progeny shown in **(f)**. Progeny from a single epimutant donor cross share the same color as their parent, while progeny from a double epimutant donor cross display mixed colors from both parents (e.g., purple for red and blue parents).

Genome-wide H3K9me remodeling was explored to determine if heterochromatin changes beyond the *fkbA* locus might contribute to FK506 and rapamycin resistance observed in the epimutants. Fourteen genes were embedded in newly gained H3K9me regions in a parent-progeny epimutant pair compared to the wildtype control (Extended Data Fig. 8c, gained H3K9me2, and Supplementary Data 4), but these were not associated with any significant transcriptional changes (Extended Data Fig. 8d and Supplementary Data 5). Similarly, genes that lost H3K9me in epimutants did not show significant expression changes (Extended Data Fig. 8e and Supplementary Data 5). Collectively, these findings indicate that the observed drug resistance is not driven by heterochromatin remodeling in this fungus. Instead, *Mucor* epimutations act posttranscriptionally through an RNAi-dependent mechanism.

These findings were generalized to other *Mucor* phylogenetic species that exhibit RNAi epimutations targeting *fkbA, M. lusitanicus* PS10 and *M. circinelloides* PS14 (6). Similarly, we found siRNAs indicative of active *fkbA* posttran-scriptional silencing, but no H3K9me2 modification at this locus and no changes in RNAP occupancy between wildtype and epimutant strains (Fig. 3c, d, PS10 and PS14, respectively). In addition to *fkbA*, previously identified epimutations conferring 5-Fluoroorotic acid (5-FOA) resistance by silencing the *pyrG* and *pyrF* pyrimidine biosynthetic genes in *M. lusitanicus* PS10 (18) were also tested by ChIP followed by quantitative PCR. Neither *pyrG* nor *pyrF* exhibited any enrichment in H3K9me2 during epimutation (Fig. 3e).

The reduction in *fkbA* transcription during epimutation, consistent with posttranscriptional gene silencing and independent of H3K9me and other chromatin modifications absent in this fungus (H3K27me and 5mC), suggests that siRNAs are functioning as exclusive epigenetic agents of inheritance. We therefore investigated whether siRNAs might be traceable from parents to their progeny. To test this, we analyzed the siRNA profiles and features in a representative set of epimutant parents and their progeny (Fig. 3f). All drugresistant epimutant progeny harbored abundant sRNAs targeting *fkbA*, which displayed features consistent with canonical siRNAs. These features include being mainly antisense to the transcript exons, having a defined length of 21 to 24 nt, and possessing uracil at the 5’ position (Fig. 3f and Extended Data Fig. 9a-c). There is also a notable bias in accumulating siRNAs towards the last exon of the *fkbA* gene.

But more importantly, a striking similarity was observed in the accumulation pattern and relative abundance of siRNAs between epimutant parents and their epimutant progeny (Fig. 3f and Extended Data Fig. 9c). The progeny from two epimutant parents displayed a blend of both parental siRNA profiles, usually favoring one parent. To validate these similarities, Principal Component Analysis (PCA) was conducted on the relative abundances of siRNA molecules complementary to the *fkbA* transcript from each epimutant. Progeny from a single epimutant donor closely clustered with their epimutant parent (Fig. 3g, red, blue, yellow, and black), whereas progeny from two epimutants were positioned between the parents, or aligned more closely to one parent (Fig. 3g, progeny in purple, red and blue parents). Taken together, these findings suggest that individual epimutants exhibit a detectable pattern of siRNA, similar to a fingerprint.

The stability of resistance in the progeny was assessed by comparing reversion dynamics and quantifying sRNA profiles in epimutant parents and progeny during subsequent mitotic passaging. As with their epimutant parents, FK506 and rapamycin resistance was maintained under drug selective pressure (Extended Data Fig. 10a). However, the amount of sRNAs did not correlate with the level of resistance, as measured by growth area in drug-containing medium after five passages (Extended Data Fig. 10b). In the absence of selective pressure, resistant isolates typically reverted to drug susceptibility before the tenth passage, following a sharp decline in siRNA levels after the first passage (Extended Data Fig. 10c, d). Therefore, resistance stability, as indicated by the number of passages required for reversion, does not correlate with either the extent of resistance (Extended Data Fig. 10e) or the sRNA levels in resistant isolates (Extended Data Fig. 10f).

## Discussion

DNA is classically regarded as the molecular basis of inheritance, but substantial evidence supports the existence of an RNA world that preceded the current central dogma of molecular biology (28). In addition to catalyzing chemical reactions, RNA might have stored genetic information and transmitted it across generations. Presently, RNA viruses transmit genetic information by hijacking the synthesis machinery of their hosts, including capsid-free, vertical RNA transmission (29). But there are more examples beyond viral mechanisms, as there is also transmission of maternal RNA and proteins to oocytes, influencing early zygotic development (30), and traumatic stress can alter microRNA expression in sperm (31, 32). Our research demonstrates that spontaneous epigenetic drug resistance can be transgenerationally inherited in living organisms, relying exclusively on RNA as the information molecule.

Our current understanding of epigenetic inheritance is mostly based on chromatin modifications, which affect DNA directly, e.g. 5mC, or DNA-binding histone proteins. However, RNA-based epigenetic inheritance remains poorly understood. Nuclear-independent, engineered RNAi epimutation inheritance has been observed across generations in the nematode *Caenorhabditis elegans* (33), but it is challenging to untangle the effects of chromatin modifications because sRNAs guide heterochromatin formation in this organism (34), and many others (35).

Many fungi rely on repressive chromatin modifications – mainly H3K9me, H3K27me, and 5mC– to silence transposable elements and influence development, growth, and virulence in animal and plant pathogens, though their contribution to gene regulation is often difficult to isolate from other genetic factors (9). Thus far, epimutations as the sole cause of gene expression changes have been convincingly demonstrated in *Mucor* spp. and *Schizosaccharomyces pombe*, where epigenetically driven silencing directly alters a discrete phenotypic trait. In *S. pombe*, caffeine resistance may arise from three independent epimutations based on H3K9me heterochromatin-mediated silencing (8). Both Clr4-mediated H3K9me and the canonical RNAi pathway are required for this process, consistent with a cotranscriptional gene silencing mechanism. Foundational studies on epigenetic inheritance have shown that heterochromatin at both endogenous and ectopic loci can be transmitted through mitosis and meiosis in *S. pombe* (36–38), especially through inactivation of the histone demethylase Epe1, which mediates a rapid epigenetic resetting mechanism. However, whether these specific epimutations are inheritable following sexual reproduction has not been reported. Interestingly, caffeine induces Epe1 inactivation in *S. pombe* (8), which may complicate its contribution to epimutation stability and heritability. It is also unclear to what extent either heterochromatin or RNAi contributes, i.e., which is the initial driver of the epimutation.

*M. circinelloides* as a model of epigenetic inheritance may offer a solution to some of these limitations. Recently, we showed that RNAi is not required to maintain H3K9me-based heterochromatin at transposable elements, suggesting that these two mechanisms function independently in *M. circinelloides* and *M. lusitanicus* (26). Our findings build on this and indicate that de novo RNAi epimutations are not triggering heterochromatin formation in either epimutants or their drugresistant progeny analyzed thus far. These insights enabled us to focus specifically on RNAi inheritance, demonstrating that siRNAs can function as the sole molecular determinants that cause, maintain, and transmit RNAi epimutations, uncoupled from chromatin modifications commonly associated with epigenetic inheritance in other organisms.

RNAi inheritability is independent of mating type. Notably, it is also independent of the genetic locus targeted by the epimutation, as shown by multiple epimutant drugresistant progeny that inherited the *fkbA* allele from a naïve, drug susceptible wildtype parent. Mucoralean sexual reproduction begins with the contact of sexually determined, engorged hyphae called zygophores, which develop into progametangia. As progametangia differentiate into gametangia, they become separated from the mycelium by septae, and fuse to form a zygosporangium and zygospore (19, 39). Plasmogamy and shared cytoplasmic contribution from both gametangia, combined with our findings, suggest cytoplasmic inheritance is the most parsimonious model for siRNA transmission. Cytoplasmic inheritance would explain the higher inheritance rates observed with two epimutant parents, implying that siRNA availability at the tip of the progametangia prior to septa formation is a crucial determinant for inheritance. Previous studies have shown that the RNAi components Dcl2, Ago1, Qip1, and Rdrp2 are essential for initiating and maintaining both epimutations and canonical gene silencing (6, 14, 15). Rdrp2 functions as the primary RdRP, amplifying secondary siRNAs and thereby propagating RNAi throughout multiple mitotic divisions (40). However, the role of Rdrp2 in siRNA inheritance is challenging to ascertain as *rdrp2*Δ mutants are unable to mate and produce zygospores (41). Given that heritable sRNAs in the epimutant progeny harbor typical features of canonical siRNAs, we propose that the canonical RNAi mechanism may extend beyond vegetative growth to preserve epimutations during sexual reproduction, similarly to *C. elegans* (42, 43).

Our findings contribute to the growing evidence supporting epigenetic transgenerational inheritance, which allows organisms to adapt rapidly to environmental pressures before genetic changes can occur. Once the environmental pressure is relaxed, epimutations can readily revert, providing remarkable adaptive phenotypic plasticity. This phenomenon is particularly concerning in pathogenic microbes that maintain epigenetic modifications (9, 44, 45) and/or RNAi machinery (46, 47). In recent years, epimutations have been identified as a novel mechanism that confers fAMR (6, 8), posing a significant threat as they may evade detection due to their transient and adaptive nature. Understanding how epimutations are inherited and the mechanisms driving resistance provide solutions to the growing challenges posed by fAMR.

## Methods

### Fungal strains, culture, drug susceptibility testing, and unstable resistant isolation

Extended Data Table 2 lists the fungal strains utilized in this work. All generated strains were derived from *M. circinelloides* CBS394.68 (PS15–) and/or CBS172.27 (PS15+) (13). To conduct experiments, asexual spores were freshly harvested after 4-6 days incubation in yeast-peptone-dextrose (YPD) at room temperature. Spores were challenged with FK506 (1 *µ*g/ml) and/or rapamycin (100 ng/ml) to obtain resistant isolates, test drug susceptibility, and maintain epimutations. Resistant isolates were selected by repeatedly culturing 200 or 2,000 spores from naïve, wildtype isolates of both PS15 mating types on FK506-containing medium for five days, until sufficient resistant mycelial colonies were recovered. After selection, resistant isolates were challenged with FK506 and rapamycin for 72 hours to confirm loss of FKBP12 function in isolates exhibiting resistance to both drugs. To test resistance stability, resistant isolates were serially passaged on non-selective, drug-free YPD for 84 hours. Isolates exhibiting complete yeast-like growth on FK506 and therefore, reverting to drug susceptibility after up to ten non-selective passages were classified as unstable resistant isolates.

### Nucleic acids isolation, blotting, immunoprecipitation, and sequencing

Mycelium cultures were collected from YPD –with or without FK506 as needed–, flash froze in liquid N_2_ and ground into a powder for nucleic acid isolation. Genomic DNA samples for Illumina sequencing were prepared using Norgen Biotek Yeast/Fungi Genomic DNA Isolation Kit from overnight, broth cultures. Genomic DNA libraries were constructed with Roche KAPA HyperPrep Kit. 150-bp paired-end reads were obtained utilizing the NovaSeq X Plus sequencing system. Concurrently, *fkbA* sequences were PCR-amplified using primer pairs listed in Extended Data Table 3 and analyzed by Sanger-sequencing. Ultra-high molecular weight DNA was isolated following a phenol/chloroform-based DNA extraction method (27) from PS15– and + 6-hour pregerminated spores. DNA was ligated with barcodes (EXP-NBD104, NB10 for PS15– and NB12 for PS15+) and sequencing adapters (SQK-LSK110) to prepare ONT MinION libraries, and loaded into a single MinION flowcell (FLO-MIN106) for a 72-hour sequencing run.

Total RNA samples from 72-hour cultures in YPD agar were prepared with QIAGEN miRNeasy Mini Kit and divided into small and long RNA preparations. sRNAs were subjected to electrophoresis in urea acrylamide gels (Invitrogen), transferred, and chemically crosslinked [1-Ethyl-3-(3-dimethylaminopropyl)carbodiimide (EDC)] to neutral nylon membranes (Hybond NX, Amersham) as previously described (48). *fkbA* sense, *α*[^32^P]UTP-labelled radioactive probes were digested into small fragments using an alkaline solution and hybridized to detect *fkbA* antisense sR-NAs, using similarly prepared *5*.*8S* rRNA antisense digested probes as a loading control. Unstable resistant isolates har-boring *fkbA* antisense sRNAs were classified as epimutants. sRNA libraries were amplified using QIAseq miRNA library kit and 75-bp single-end reads were sequenced with Illumina NextSeq 1000 High-Output sequencing system. rRNA-depleted RNA libraries (long RNA) were prepared using Illumina Stranded Total RNA Prep and Ribo-Zero Plus rRNA Depletion Kit using *M. circinelloides* rDNA-specific probes, and 150-bp paired-end reads were obtained with NovaSeq X Plus sequencing system.

Overnight, YPD broth cultures were processed for ChIP as described in previous research (26, 49), preparing two biological replicates per sample. Briefly, cultures were crosslinked, lysed, and chromatin sonicated throughout 35 cycles 30 s ON/OFF in a Bioruptor (Diogenode) to obtain 100-300 bp chromatin fragments. Sheared chromatin aliquots were stored as input DNA controls. ChIP-grade antibodies *α*-H3K9me2 (ab1220, Abcam) and *α*-RNA pol II (39497, Active Motif), as well as protein A or G magnetic beads accordingly, were utilized to immunoprecipitate chromatin-bound DNA. After washing and decrosslinking, DNA was purified by a phenol/chloroform-based extraction method. DNA preparations from the same strain and conditions were pooled together. Libraries were prepared using Roche KAPA Hyper-Prep Kits and sequenced with Illumina NovaSeq X Plus sequencing system for 150-bp paired-end reads. For quantitative PCR, ChIP and Input DNA were amplified using SYBR Green PCR Master Mix (Applied Biosystems) with specific primer pairs (Extended Data Table 3). qPCRs were performed in triplicate using a QuantStudio™ 3 real-time PCR system. Fold enrichment was determined as a relative quantification to the Input DNA (2^-ΔCT^) and to the negative control *vma1* values (2^-ΔΔCT^), a constitutively active gene lacking heterochromatin marks.

### Genome assembly, annotation, and homolog identification

Long, ONT-based, FASTQ raw reads were obtained from MinION raw output and demultiplexed into PS15– and + reads with ONT basecaller Dorado v0.7.0 (https://github.com/nanoporetech/dorado/). Draft assemblies were generated using Flye v2.9.3-b1797 (50) and polished with Racon v1.5.0 (51). Assemblies were further corrected with NextPolish1 v1.4.1 (52) and 2 v0.2.0 (53) using long, ONT-based and Illumina-based, adapter-trimmed short reads. Long and short reads were aligned with Minimap2 v2.26-r1175 (54) to examine coverage as a measure of assembly correctness. Similarly, contig sequences were aligned and visualized using Circos v0.69-8 (55) to examine synteny between both assemblies (Extended Data Fig. 4). 13 alignment blocks that showed perfect synteny and contiguity between both assemblies –referred to as synteny blocks– were selected for further analyses. Repeated sequences were identified utilizing RepeatModeler2 v2.0.5 (56) and classified by iterative similarity searches against the RepBase database for RepeatMasker (Edition-20181026) (57) and already classified repeats. After masking the genomes with these repeat libraries and RepeatMasker v4.1.6 (http://www.repeatmasker.org/), gene annotation was conducted using the Funannotate pipeline v1.8.15 (58) and training models generated with our RNA data. Genome assemblies were evaluated using QUAST v5.2.0 (59) for contiguity metrics and the eukaryotic ortholog database (eukaryota_odb10) from BUSCO v5.7.1 (60) for completeness (Extended Data Table 1). RNAi components were identified by NCBI BLASTp v1.12.0 (61) searches using *M*.*lusitanicus* protein sequences as queries, extracted from M. *lusitanicus* MU402 v1.0 available at the Joint Genome Institute Mycocosm platform (https://mycocosm.jgi.doe.gov/). Identified proteins that also returned positive reciprocal BLASTp hits were retained, and their protein domain configuration was predicted with InterProScan v5.70-102.0 (62) to evaluate their putative function (Extended Data Fig. 1a).

### Mating assays, zygospore isolation, and progeny germination

Opposite mating-type spores were inoculated onto four spots in whey agar plates, allowing them to grow facing each other in dark conditions and room temperature. After 7 days, zygospores that formed at the intersection of opposite mating types were scrapped from the agar using tweezers and suspended in distilled water. Zygospores were germinated according to previous research (13, 20, 22). Briefly, the suspensions were successively pass through 40 and 100 µm cell strainers, discarding asexual spores and hyphal fragments and therefore, enriching in zygospores. Single zygospores were manually dissected onto agarose 1% plates at pH 4.0 and incubated at room temperature for 2-8 weeks until germi-nation. Germspores from germinating zygospores were collected in water using a wet micropipette tip.

### Genetic variation analyses

Low quality and adapter sequences were removed from genomic DNA raw reads using fastp v0.23.4 (63). For this and subsequent analyses, unless otherwise stated, the newly generated PS15-genome was selected as the reference genome to directly compare all of the samples. Processed reads were aligned with BWA-MEM v0.7.17-r1188 (64) ensuring Picard compatibility, and resulting aligments were sorted and further processed to mark duplicates and assigned read groups using the GATK4 suite v4.4.0.0 (65).

For meiotic recombination analyses, variant calling was performed on PS15– and + by GATK4 HaplotypeCaller. Low quality variants (QD < 20.0, QUAL < 30.0, SOR > 3.0, FS > 60.0, MQ < 40.0) and variants that were concordant between PS15– and PS15+ were filtered by GATK4 VariantFiltration, as well as variants that overlapped with repeated sequences using Bedtools v2.30.0 (66), generating high-quality, PS15+ specific SNVs derived from non-repetitive genomic sequences. A total of 18,731 SNVs overlapped with the synteny blocks previously identified and were used to characterize genetic variation in the progeny datasets. Parent and progeny processed alignments were piled up against this set of variants, establishing a threshold of *≥*95% read support towards either reference (PS15–) or alternate (PS15+) position to be considered inherited SNVs. Same parent variants closer than 50 kb were merged to avoid overplotting during rendering (Extended Data Fig. 5).

To identify potential mutations conferring FK506 and rapamycin resistance, a more stringent variant calling analysis was applied to the processed alignments from parental samples (PS15–, E1– to E6–, PS15+, E10+, E12+, and E15+). Variants were called using GATK4 HaplotypeCaller and filtered as described above, but with stricter thresholds (QD < 5.0, DP < 60.0, QUAL < 300.0, SOR > 1.0, FS > 0.1, MQ < 60.0). Variants failing these filters were excluded along those shared with either wildtype control (PS15– or PS15+) and those overlapping repetitive sequences. The resulting high-confidence variants were manually curated by examining the processed alignment from their corresponding wild type; variants already present in the wildtype reads were considered false positives and discarded (Extended Data Fig. 7). DNA coverage was assessed with Deeptools2 bamCoverage v3.5.4 (67) and coverage plots were centered to approximately whole-genome average coverage, which was assumed to represent a haploid genome (Extended Data Fig. 6).

### RNA-sequencing analyses

sRNA raw reads were trimmed by Trim Galore! v0.6.10 (https://github.com/FelixKrueger/TrimGalore) and stringent parameters (−-stringency 4) to avoid random 3’-end trimming. Processed reads were aligned using Bowtie v1.3.1 (68). ShortStack v3.8.5 ^69^ was run on the resulting alignment files to identify sRNA-producing loci agnostically. Bases of sRNA-producing loci overlapping coding, intergenic, and repetitive regions were quantified and visualized as a pie chart (Extended Data Fig. 2a). Additionally, sRNAs mapping to annotated gene features were quantified using ShortStack as previously. Coverage files were generated using bamCoverage and normalized to counts per million (CPM).

Alternatively, processed reads were aligned with STAR v2.7.11b (69) to assess splice junction read support (Extended Data Fig. 9c). Samtools v1.10 (70) and Bioawk v20110810 (https://github.com/lh3/bioawk) were used to count and/or filter reads harboring specific siRNA features, namely antisense orientation, length of 21-24 nt, and 5’-U. Relative abundances of siRNA reads were computed (read number *\* total fkbA reads), and this standardized relative abundances were utilized to perform a PCA to assess similarities among samples.

Long RNA raw reads were trimmed with fastp and aligned with STAR. Alignments were split into forward and reverse strand to generate stranded coverage files with bamCoverage and normalized to CPM. Differential expression was then quantified using DESeq2 v1.44.0 (71), followed by a log_2_ transformation of CPM. Expression differences were considered significant at an adjusted p-value or False Discovery Rate below *≤*0.05 (FDR 0.05). Normalized expression values were visualized in a heatmap generated with ComplexHeatmap v2.20.0 (72) (Extended Data Fig. 8d, e).

### Chromatin immunoprecipitation-sequencing analyses

Raw read samples were processed with fastp to remove adapter and low-quality sequences. Processed reads were aligned with BWA-MEM. Coverage of ChIP-enrichment was determined as the IP/Input DNA ratio using bamCompare v3.5.4 and normalized to CPM. Broad ChIP-enriched regions were identified by MACS2 v2.2.9.1 (73). Differential binding analysis was performed using DiffBind 3.14.0 (74), including read quantification and normalization, contrast definition and identification of RNA polymerase II binding differences between epimutant and wildtype samples, specifically at the *fkbA* locus (Extended Data Fig. 8a).

Additionally, genes with *≥* 90% of their sequence embedded in H3K9me-based heterochromatin were identified. Shared H3K9me2-embedded genes between epimutant and wildtype samples were visualized as a Venn diagram generated by ggVennDiagram v1.5.2 (75) (Extended Data Fig. 8c).

### Statistical information

Inheritance ratios in the progeny were compared to the expected values of monogenic trait inheritance in haploid organisms with only one parent exhibiting the trait (1:1 ratio, *p* = 0.5, 0.5), or both parents exhibiting the trait (1:0 ratio, *p* = 0.95, 0.05 to avoid expected frequency = 0 and to account for other biological variation, e.g., spontaneous mutations). Statistical significance was assessed by goodness-of-fit (GOF) Chi-square tests (1 degree of freedom) with a significance level of 0.05.

One-way ANOVA testing was conducted to determine significant differences among ChIP qPCR fold enrichment values (n = 3), using Tukey’s Honestly Significant Distance (HSD) to avoid false positives due to multiple pairwise comparisons. Tests were performed with a 95% CI and pairwise comparison results were summarized by a compact letter display using the R package multcomp (76).

## Data availability

Genome assemblies, gene annotation, and raw ONT and Illumina-based raw FASTQ reads are publicly available under NCBI’s Sequence Read Archive (SRA) project accessions PRJNA1168935 (PS15–) and PRJNA1168941 (PS15+). The remaining raw sequencing data can be accessed under PRJNA1170303, including small RNA, rRNA-depleted RNA, ChIP, and Illumina wholegenome sequencing. In addition, *M. lusitanicus* PS10 wildtype ChIP and small RNA data was retrieved from the publicly available project accession PRJNA903107 and used to compare our findings. Individual SRA run (SSR) accession numbers are listed in Supplementary Data 6. Raw data underlying bar and scatter plots, as well as uncropped and unprocessed scans of Northern blots is provided as multiple labeled files within a zip-compressed Source Data file.

## Author contributions

C.P.-A, M.I.N.-M, and J.H. conceptualized the project; C.P.-A and M.I.N.-M designed the experiments; C.P.-A, M.I.N.-M, and Z.X. performed the experiments; G.W. provided biological resources and critical advice on mating; C.P.-A and M.I.N.-M assembled and annotated the genomes; C.P.-A and M.I.N.-M analyzed and interpreted the data; C.P.-A and M.I.N.-M generated figures; C.P.-A and M.I.N.-M wrote the original draft; C.P.-A, M.I.N.-M, and J.H. edited the original draft; C.P.-A, M.I.N.-M, Z.X., J.H., and G.W. reviewed the final manuscript.

## ACKNOWLEDGEMENTS

We thank our lab manager Anna Floyd Averette for her support. We commend Dr. Devjanee Swain Lenz and Duke’s Sequencing and Genomic Technologies Core Facility for their assistance, as well as Thomas Milledge and the Duke Computer Cluster team for their computing resources. We thank Prof. Thomas Petes, Prof. Francisco E. Nicolás, Dr. Sheng Sun, and Dr. Jun Huang for critical reading and insightful suggestions on the manuscript. This study was supported by NIH/National Institute of Allergy and Infectious Diseases grants R01-AI170543, R01-AI39115, and R01-AI050113 awarded to J.H. The funders had no role in study design, data collection and interpretation, or the decision to submit the work for publication. J.H. is Codirector and Fellow of the CIFAR program Fungal Kingdom: Threats & Opportunities.

## Supplementary information

**Extended Data Table 1.**
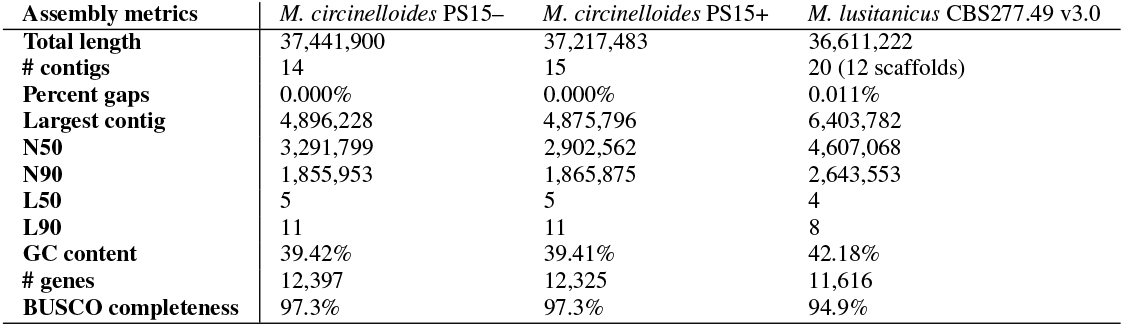
Genome assembly metrics for *M. circinelloides* PS15 opposite mating types. Values are compared to the latest *M. lusitanicus* CBS277.49 genome assembly. Assembly completeness is assessed as the percentage of conserved orthologs from the BUSCO Eukaryota ortholog database (eukaryota_odb10).

**Extended Data Table 2.**
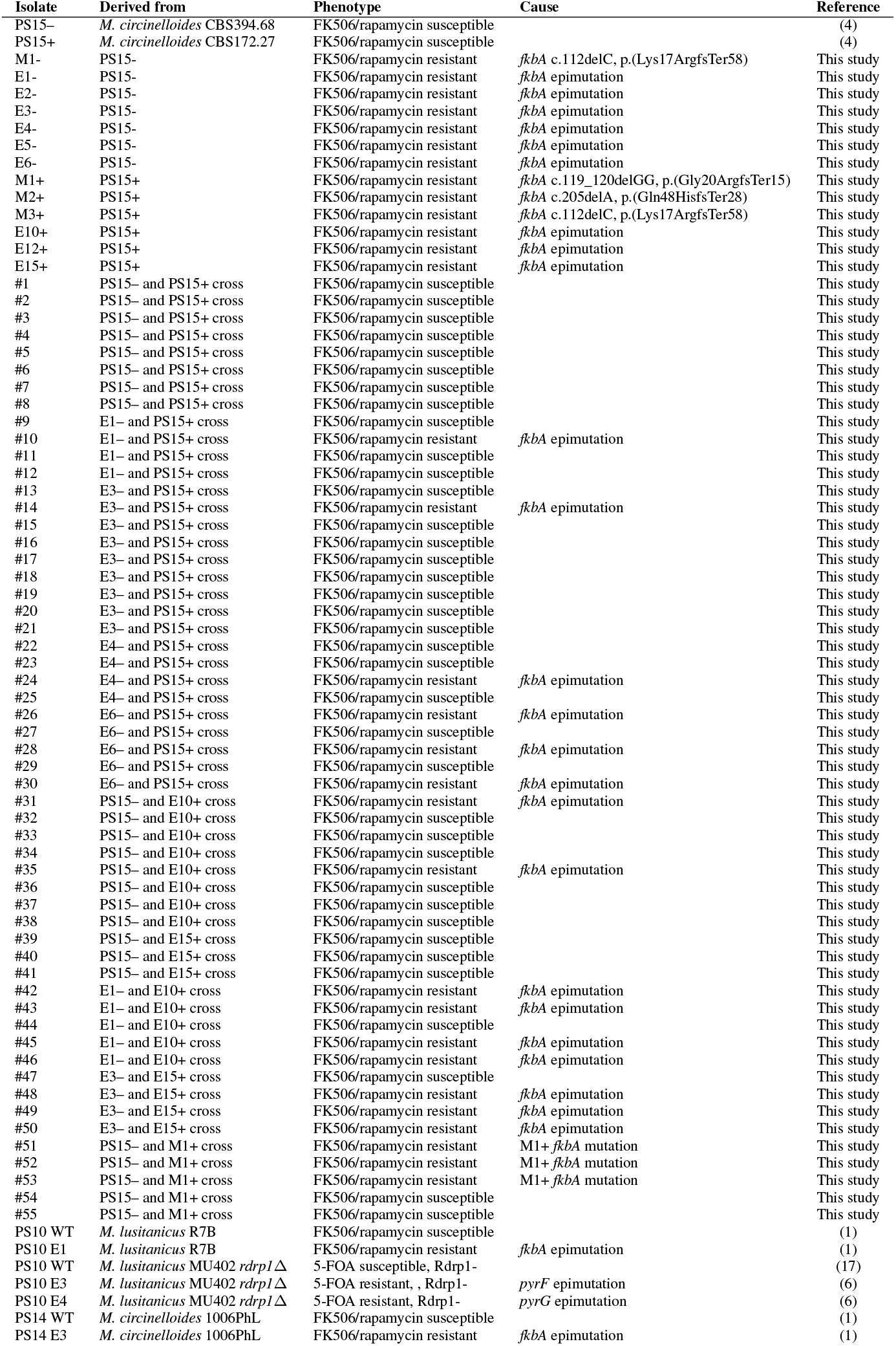
Strains utilized in this work are listed highlighting the isolate name in the manuscript, their parental strains, relevant phenotype, cause of drug resistance, and reference.

**Extended Data Table 3.**
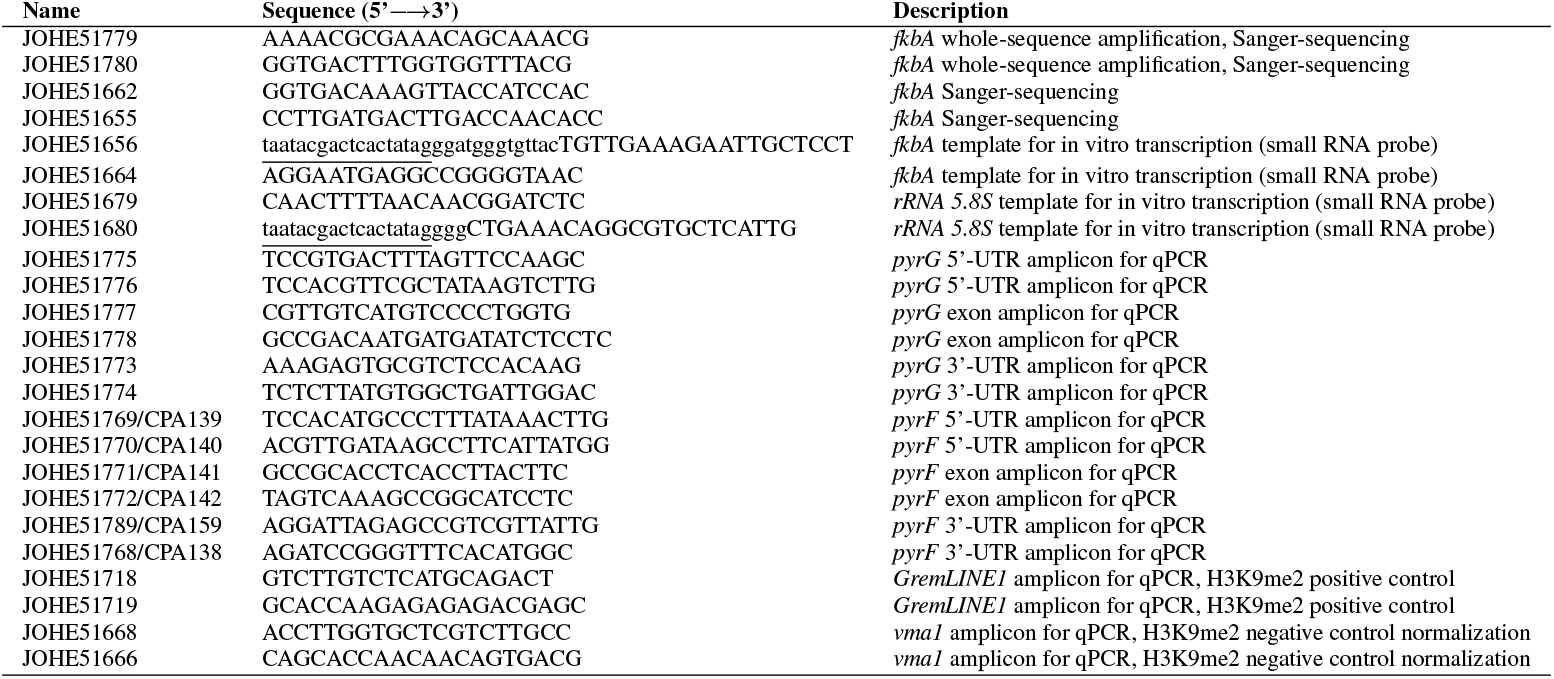
Primers used for this study are listed including their sequence, and brief description. Lowercase characters indicate primer tails containing the T7 promoter sequence (underlined) and a short spacer sequence for in vitro riboprobe transcription.

**Extended Data Fig. 1.**
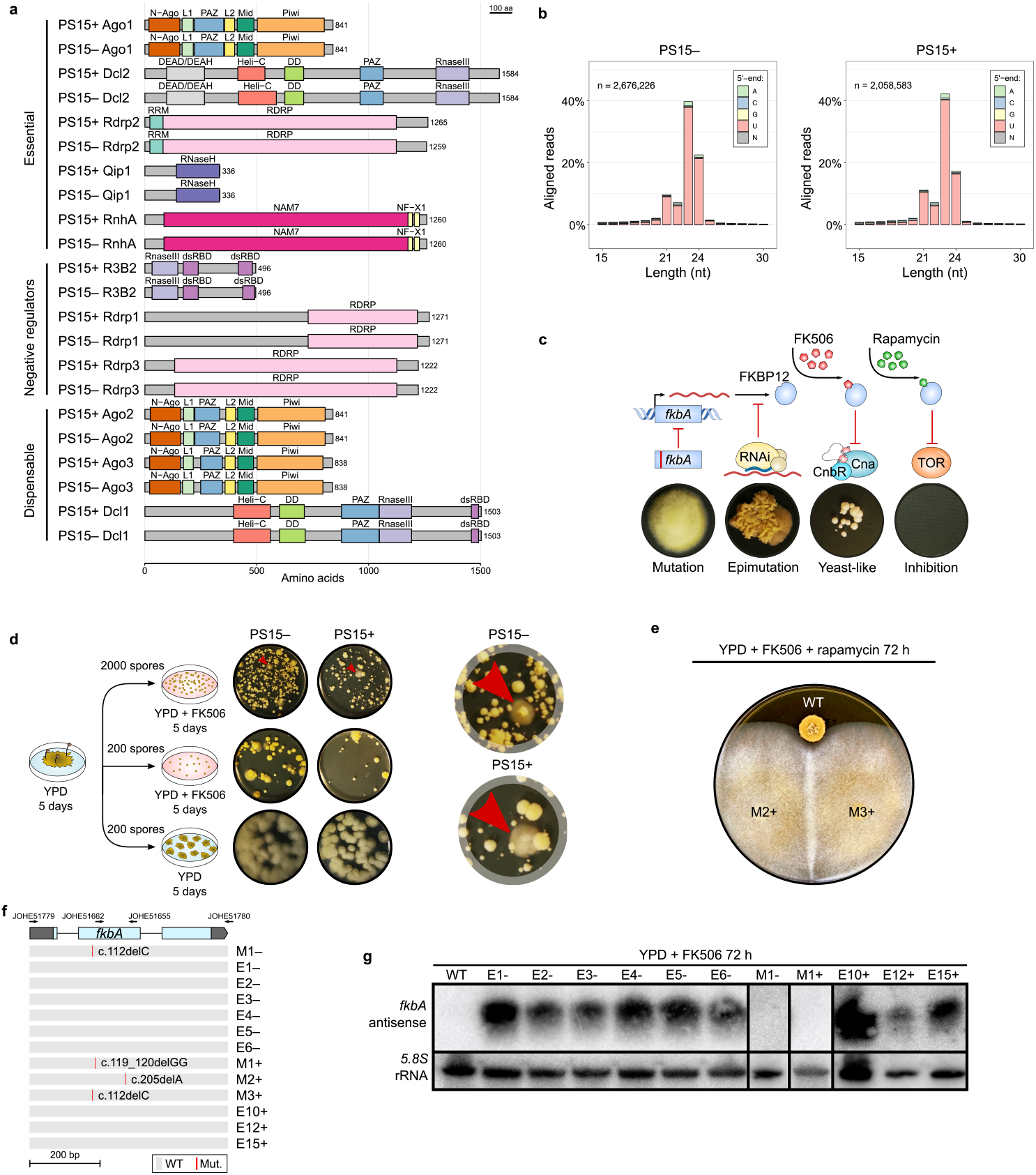
*Mucor circinelloides* PS15 RNAi proficiency enables *fkbA* silencing. **(a)** The RNAi components are classified into essential, negative regulators, and dispensable for epimutation. Rectangles display the full-length, scaled protein sequences of every identified protein homolog and their predicted, color-coded InterPro protein domains abbreviated as follows: N-Ago (Protein argonaute, N-terminal, IPR032474); L1 (Argonaute, linker 1 domain, IPR014811); PAZ (PAZ domain, IPR003100); L2 (Argonaute, linker 2 domain, IPR032472); Mid (Protein argonaute, Mid domain, IPR032473); Piwi (Piwi domain, IPR003165); DEAD/DEAH (DEAD/DEAH box helicase domain, IPR011545); Heli_C (Helicase C-terminal domain-like, IPR001650); DD (Dicer dimerization domain, IPR005034); RNaseIII (Ribonuclease III domain, IPR000999); RRM (RNA recognition motif domain, IPR000504); RDRP (RNA-dependent RNA polymerase, eukaryotic type, IPR007855); RNaseH (Ribonuclease H domain, IPR012337); NAM7 (DNA2/NAM7-like helicase, IPR045055); NF-X1 (Zinc finger, NF-X1-type, IPR000967); dsRBD (Double-stranded RNA-binding domain, IPR014720). **(b)** The percentage of sRNA aligned reads is plotted according to length and 5’-end nucleotide distribution (color-coded). **(c)** The mechanism of action of FK506 and rapamycin via FKBP12 binding and inhibition of calcineurin (Cna and CnbR) or TOR is illustrated. *fkbA* mutations (red line) or RNAi epimutations cause loss of FKBP12 function and result in drug resistance. **(d)** Isolates exhibiting loss of FKB12 function were obtained by exposing 200 or 2,000 spores to FK506 and screening for mycelium formation as an indication of resistance. Red arrows highlight representative resistant colonies, shown in the zoomed-in view on the right. **(e)** <u>M</u>utant (M) isolates M2+ and M3+ exhibiting FK506 resistance were challenged with both FK506 and rapamycin for 72 hours, comparing growth to wildtype strains (WT). Isolates resistant to both drugs are labelled in black, and susceptible isolates in white. **(f)** Schematic representation of the *fkbA* gene, showing 5’ and 3’ untranslated regions (UTRs, dark gray blocks) and coding regions (light blue blocks). Primers used for Sanger sequencing are indicated above the gene diagram. Below, the *fkbA* nucleotide sequence is shown, with identified mutations highlighted in red as opposed to gray for wildtype sequences. **(g)** Small RNA Northern blot of depicted isolates after a 72-hour exposure to FK506, probed for antisense *fkbA* small RNAs (*fkbA* antisense, above) and sense *5*.*8S* ribosomal RNA (*5*.*8S* rRNA, below) as a loading control.

**Extended Data Fig. 2.**
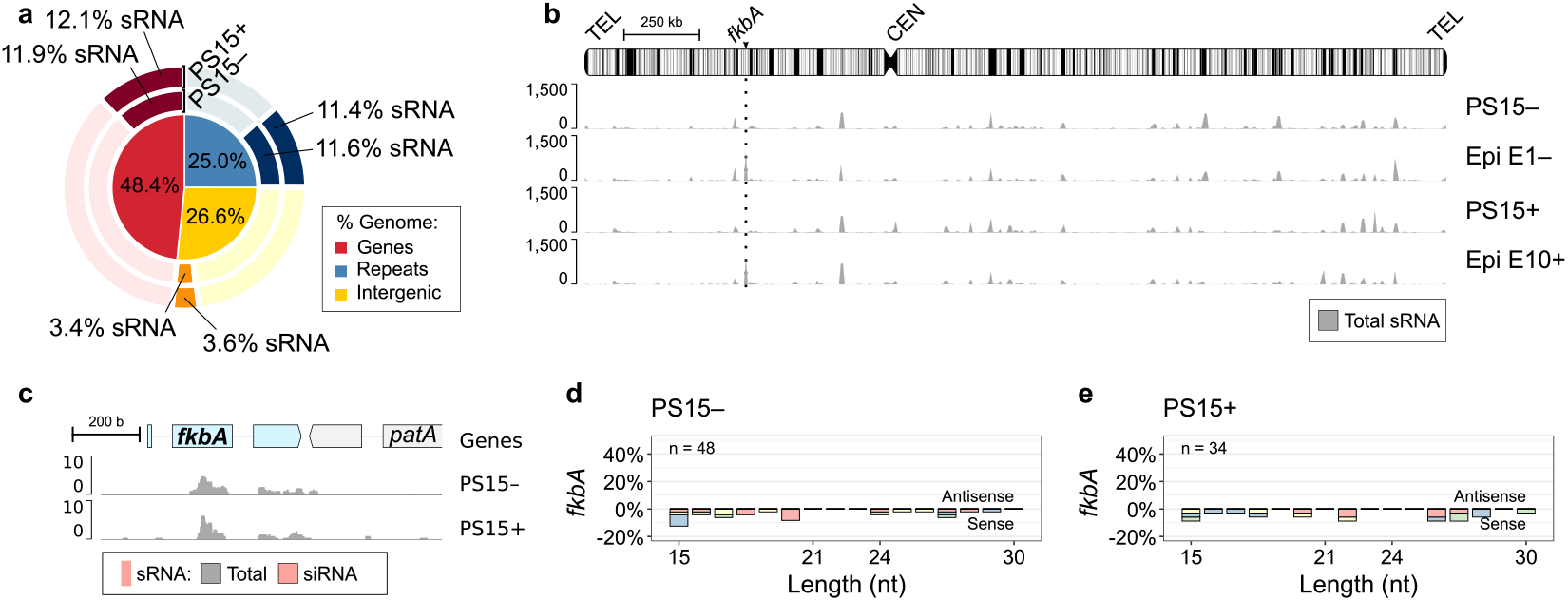
Wildtype PS15 opposite mating types are RNAi-proficient but do not silence the *fkbA* gene in the absence of selection. **(a)** Proportion of gene, repetitive, and intergenic sequences in the PS15^−^ genome (inner pie chart, color-coded). Outer pie chart shows the corresponding proportions of base pairs overlapped by sRNA loci (darker shade) and not overlapped (lighter shade). **(b)** Genome-wide distribution of total sRNA (dark gray) across the chromosome carrying *fkbA* (arrowhead and dotted line) in wildtype, naive PS15– and PS15+, as well as representative epimutants of each mating type. Repeats are shown as black vertical lines; the centromere (CEN) as a constriction; telomeres (TEL) as rounded ends. **(c)** sRNA coverage (dark gray) and siRNA (red) across the *fkbA* locus in PS15– and PS15+. **(d, e)** Length distributionand 5’-end nucleotide composition (color-coded) according to strand sense of sRNA reads mapping to *fkbA* in wildtype, naive PS15– (d) and PS15+ (e).

**Extended Data Fig. 3.**
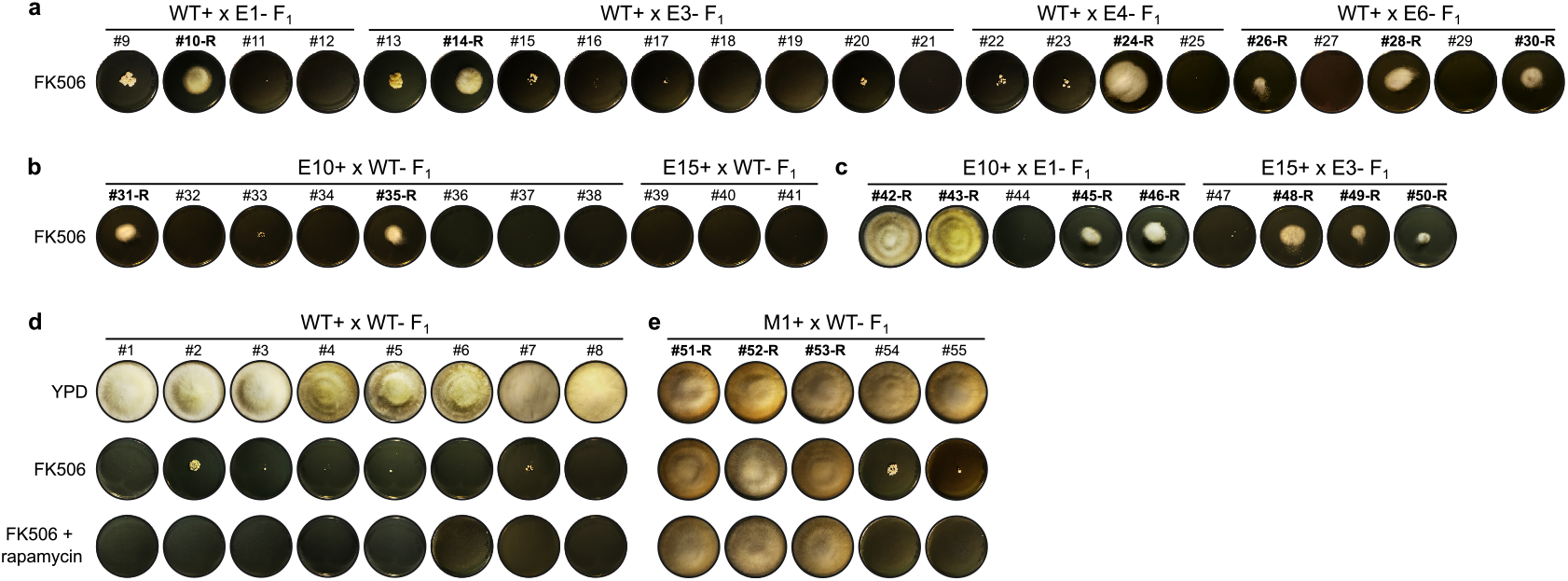
Independent FK506 challenge and F_1_ progeny derived from control crosses. **(a-c)** FK506 susceptibility was assessed in the progeny from **(a)** – mating type epimutant crossed with + mating type wildtype strain; **(b)** + mating type epimutant crossed with – mating type wildtype strain; and **(c)** a cross between two epimutants (– and +). This independent challenge with FK506 revealed the same susceptibility patterns as shown in Fig. 2a-c for FK506 and rapamycin combined exposure, indicating the observed resistance was already present in the germspores before the drug challenge, as opposed to arising de novo due to treatment. **(d, e)** Susceptibility to FK506 alone and in combination with rapamycin was determined in the progeny from **(d)** a cross between two wild type (– and +); and **(e)** – mating type *fkbA* mutant crossed with + mating type wildtype strain. Growth in YPD served as a viability control.

**Extended Data Fig. 4.**
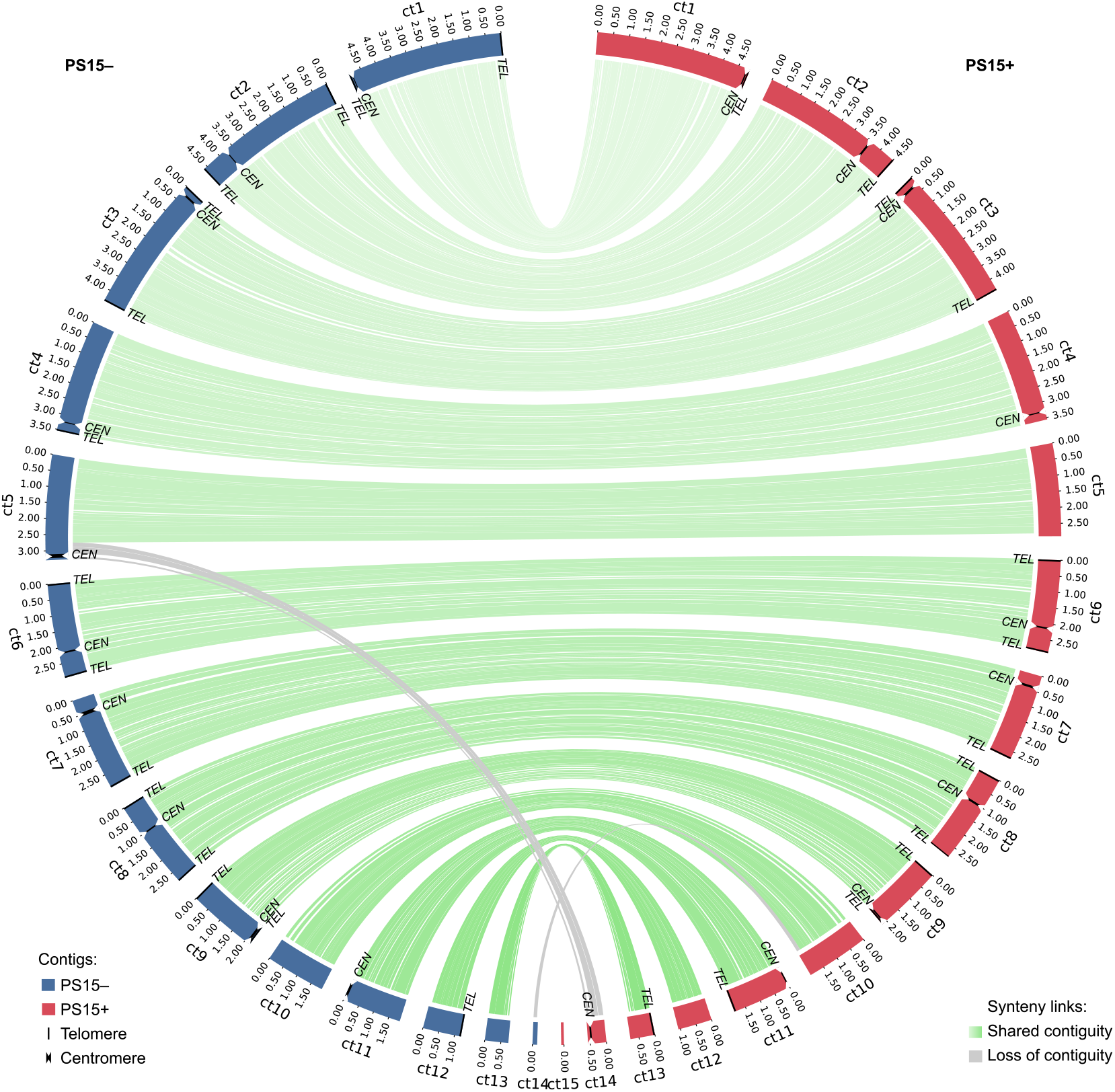
*Mucor circinelloides* PS15 – and + genome assemblies. Contig distribution and length of each assembly are shown as colored rectangles (– in blue, + in red). Centromeres (*CEN*) are depicted as constrictions and telomeres (*TEL*) are indicated by black lines closing the contig. Synteny between homologous contigs from each strain is shown as green lines. In total, 13 shared synteny blocks were defined and used in meiotic recombination and coverage analyses shown in Extended Data Fig. 5 and 6, comprising six complete telomere-to-telomere chromosomes (ct1, ct2, ct3, ct6, ct8, and ct9) and seven contigs (ct4, ct5, ct7, ct10, ct11, ct12, ct13), as well as nine centromeres located on ct1, ct2, ct3, ct4, ct6, ct7, ct8, ct9, and ct11. On the other hand, gray lines indicate similarity between regions that are not contiguous in both strains and were excluded from further meiotic recombination and coverage analyses. One centromere lies outside the defined synteny blocks. In PS15–, it is located on ct5, whereas in PS15+, the homologous region is split between ct14 (which harbors the centromere) and ct5. Due to unresolved chromosomal linkage between ct14 and ct5 in PS15+, the centromere-containing region was excluded from the synteny blocks used in recombination analyses.

**Extended Data Fig. 5.**
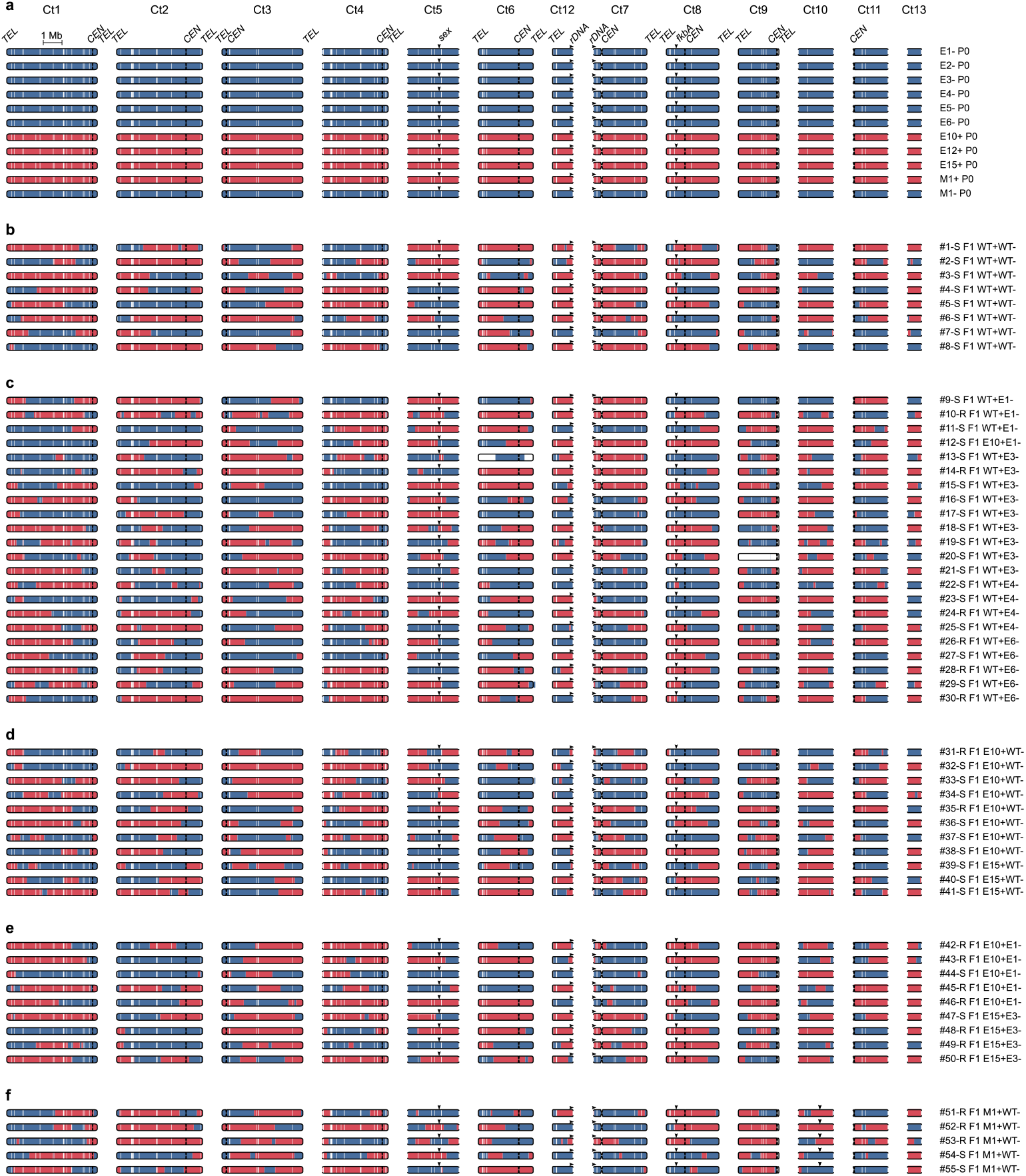
Meiotic recombination sites in F_1_ progeny. **(a-f)** Contigs exhibiting synteny and contiguity in both PS15– and + are shown to indicate meiotic recombination sites in: **(a)** parental strains (P_0_) utilized for sexual crosses, and F_1_ progeny derived from **(b)** a cross between two wild type (– and +); **(c)** – mating type epimutant crossed with + mating type wildtype strain; **(d)** + mating type epimutant crossed with – mating type wildtype strain; **(e)** a cross between two epimutants (– and +); and **(f)** – mating type *fkbA* mutant crossed with + mating type wildtype strain. The contig (Ct) drawings depict the inheritance pattern of genetic variation identified in both opposite mating types (SNPs, – mating type in blue and + mating type in red). Incomplete chromosomes, i.e., without telomeric repeats are depicted as rounded rectangles with open ends, whereas full chromosomes show a centromere (constriction) and telomeric repeats on both ends (closed ends). Relevant loci are designated by arrows (RFLPs, rDNA, *sex*, and *fkbA*), indicating their direction (forward or reverse) when relevant (rDNA). To facilitate visualization, Ct12 is shown horizontally inverted and placed to the left of Ct7 to suggest that they are most likely joined at the rDNA repeats, as both contig ends show genetic linkage in all progeny.

**Extended Data Fig. 6.**
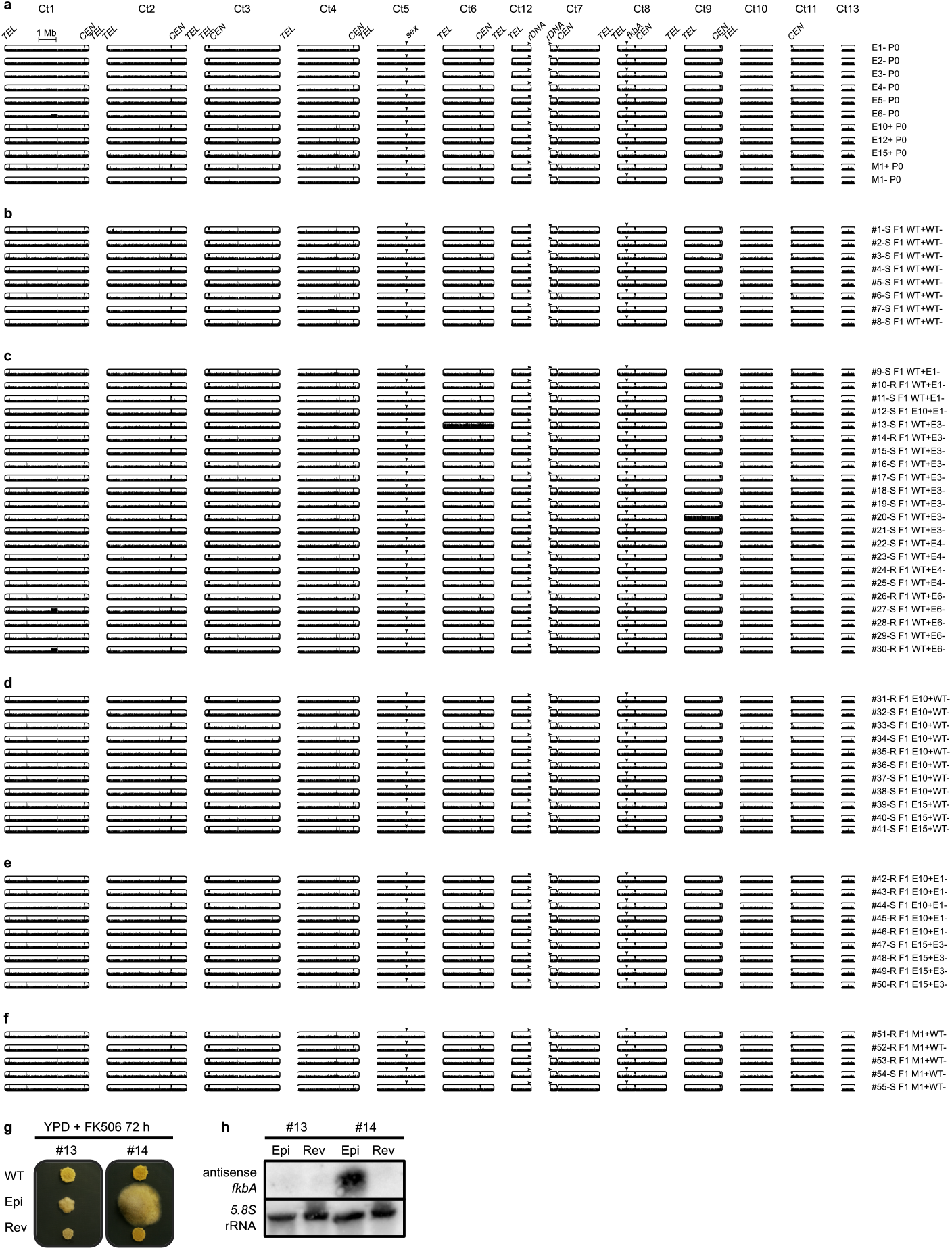
Changes in ploidy or segmental duplications or deletions are not linked to RNAi epimutations. **(a-f)** Isolates, contigs, and their features are arranged as indicated in Extended Data Fig. 5. To facilitate visualization of changes in ploidy, DNA coverage was normalized as counts per million. An E6-subpopulation harbored a segmental duplication in contig 1 (Ct1), which was inherited by two progeny (isolates #27 and #30). This duplication was unlinked to changes in FK506 susceptibility. In addition, progeny isolates #13 and #20 showed chromosome 6 (Ct6) and 9 (Ct9) duplication, respectively. Ct9 duplication does not result in FK506 and/or rapamycin resistance. **(g)** Progeny #13 (tolerant, Ct6 duplication) and #14 (resistant, euploid) were passaged on non-selective medium for ten 72-hour passages (revertants) and transferred onto FK506-selective medium next to their corresponding wildtype susceptible control and resistant isolates to assess reversion. **(h)** Small RNA Northern blot of progeny isolates #13 and #14 after a 72-hour exposition to FK506. The blot shows probe hybridizations to detect antisense *fkbA* small RNAs (*fkbA* antisense, above) and sense *5*.*8S* ribosomal RNA (*5*.*8S* rRNA, below) as a loading control. Isolate #13 did not harbor sRNAs. Although chromosome 6 duplication may confer tolerance to FK506 and rapamycin, tolerant isolate #13 exhibits yeast-like morphology and lacks antisense sRNAs targeting *fkbA*.

**Extended Data Fig. 7.**
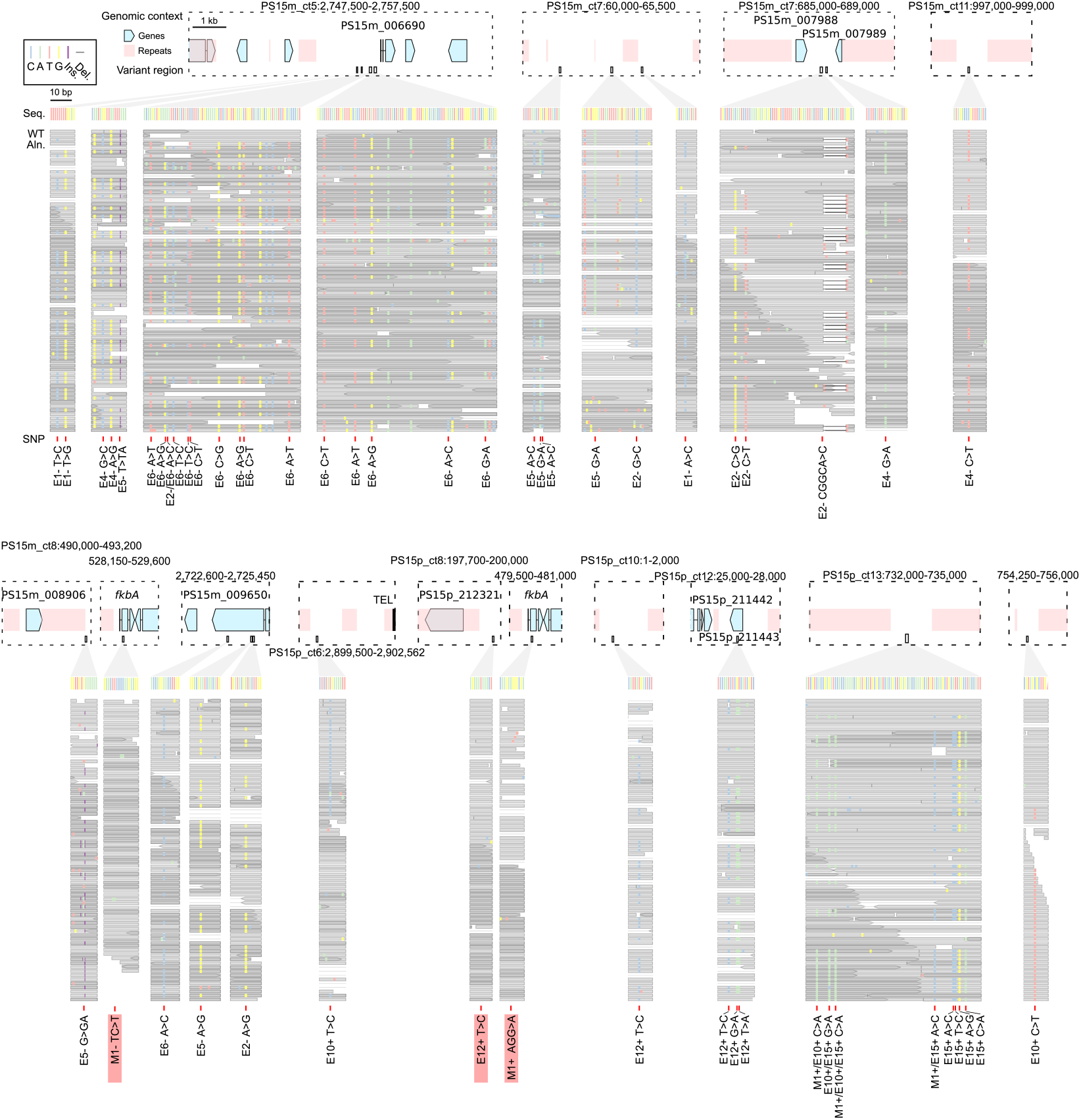
Whole-genome sequencing reveals no consistent mutations conferring FK506 and rapamycin resistance beyond the RNA-silenced, epimutant *fkbA* locus. Putative single nucleotide polymorphisms (SNPs, red) are shown within their genomic context (top; coordinates indicated). Genes are represented as light blue arrowed blocks, repeats as light red blocks, and a telomeric region (TEL) as a black block. Genes nearest to each variant are labeled. Below, zoomed-in panels (corresponding to boxed regions above) display the reference sequence (Seq.) and wildtype read alignments (WT Aln.), both color-coded as indicated. Wildtype read alignments show pre-existing variants to assess whether putative SNPs are false positives or bona fide mutations. Variants not reproduced in the wildtype alignment reads, regarded as bona fide mutations, are highlighted with red shading.

**Extended Data Fig. 8.**
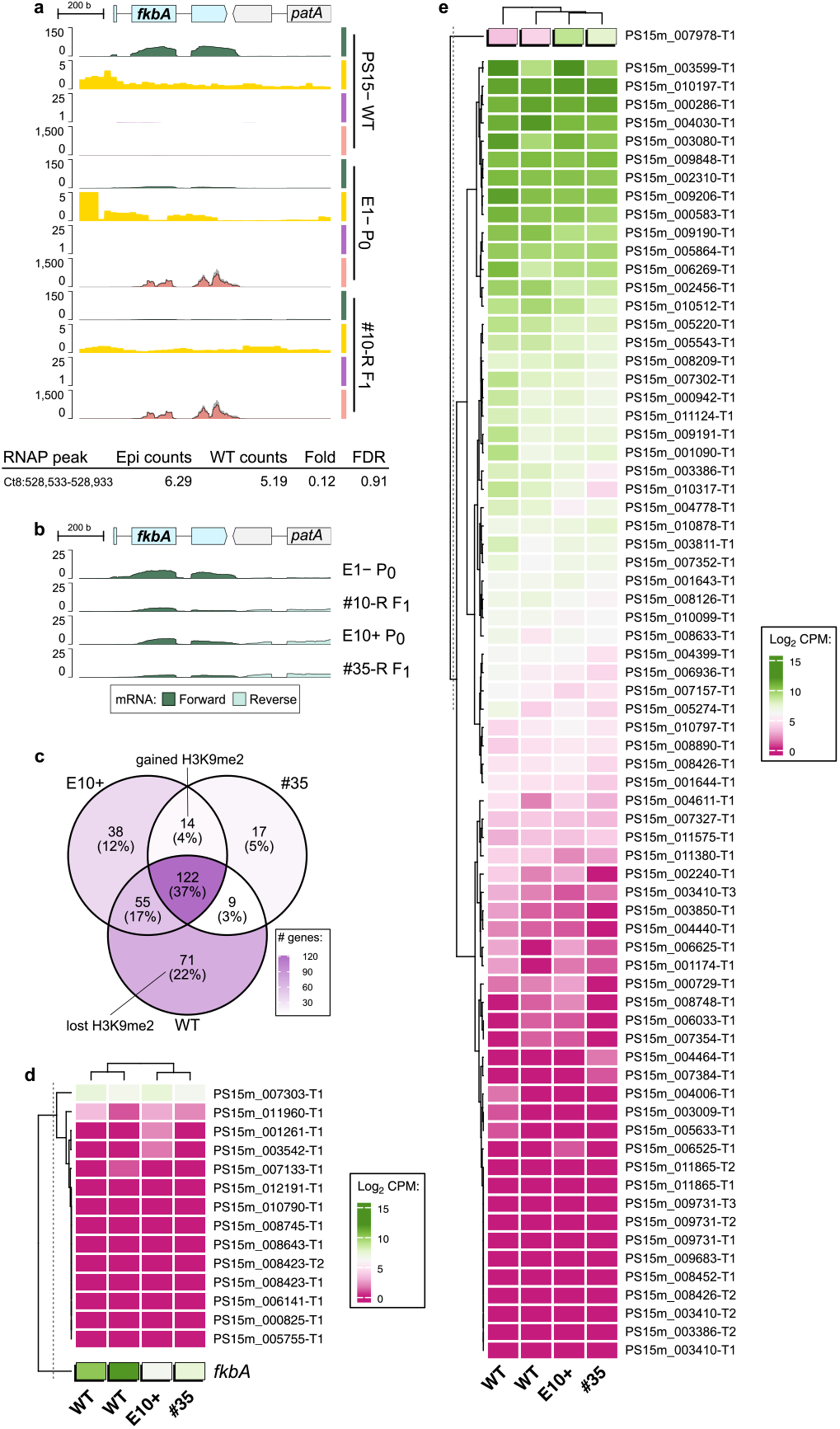
Epimutations are consistent with posttranscriptional gene silencing. **(a)** Genomic tracks showing H3K9me2 enrichment (purple), RNA Polymerase II enrichment (RNAP, yellow), stranded mRNA (green; forward and reverse overlaid), and sRNA (dark gray for total sRNA, overlaid with red for siRNA) in *M. circinelloides* PS15– wildtype, the epimutant parent (E1– P0), and its F1 progeny (#10-R F1) across an expanded view of the *fkbA* locus. Below, differential binding analysis (DiffBind, DESeq2) reveals no significant differences in RNAP enrichment between epimutant and wildtype isolates (Fold enrichment = 0.12, FDR = 0.91). **(b)** Stranded mRNA across the *fkbA* locus displayed at a lower CPM range to highlight subtle *fkbA* expression in epimutants and their progeny from both mating types. **(c)** A Venn diagram shows overlapping genes embedded in H3K9me2 (*≥* 0.05 90% of their coding sequence) in a naïve wild-type strain (WT), the epimutant E10+, and its progeny (#34). Color-coded shades of purple indicate the number of overlapping genes. Gained and lost H3K9me2 genes are highlighted, defined as those embedded in both epimutant and progeny but absent in WT (gained), or present in WT only (lost). **(b, c)** The heatmaps display color-coded gene expression values as normalized log_2_ CPM across two wildtype duplicates, an epimutant (E10+), and its progeny (#34), which served as duplicates in the differential expression analysis (DeSEQ2). Significant expression changes (FDR *≤* 0.05) between wild type and epimutant are indicated by raised, black-shadowed cells. The similarity between gene expression values and samples is depicted by rows and columns dendrograms generated through complete clustering using the Euclidean distance matrix. A dashed vertical line indicates a split in the dendrogram node, which was used to mark the positive control gene. In **(d)**, fourteen H3K9me2 gained genes are displayed, using *fkbA* as a positive control for expected expression values due to gene silencing. In **(e)**, 71 H3K9me2 lost genes are shown, using *PS15m_007978* as a positive control of an upregulated gene. Except for the positive controls, none of these genes showed significant expression changes as a consequence of gaining or losing H3K9me2.

**Extended Data Fig. 9.**
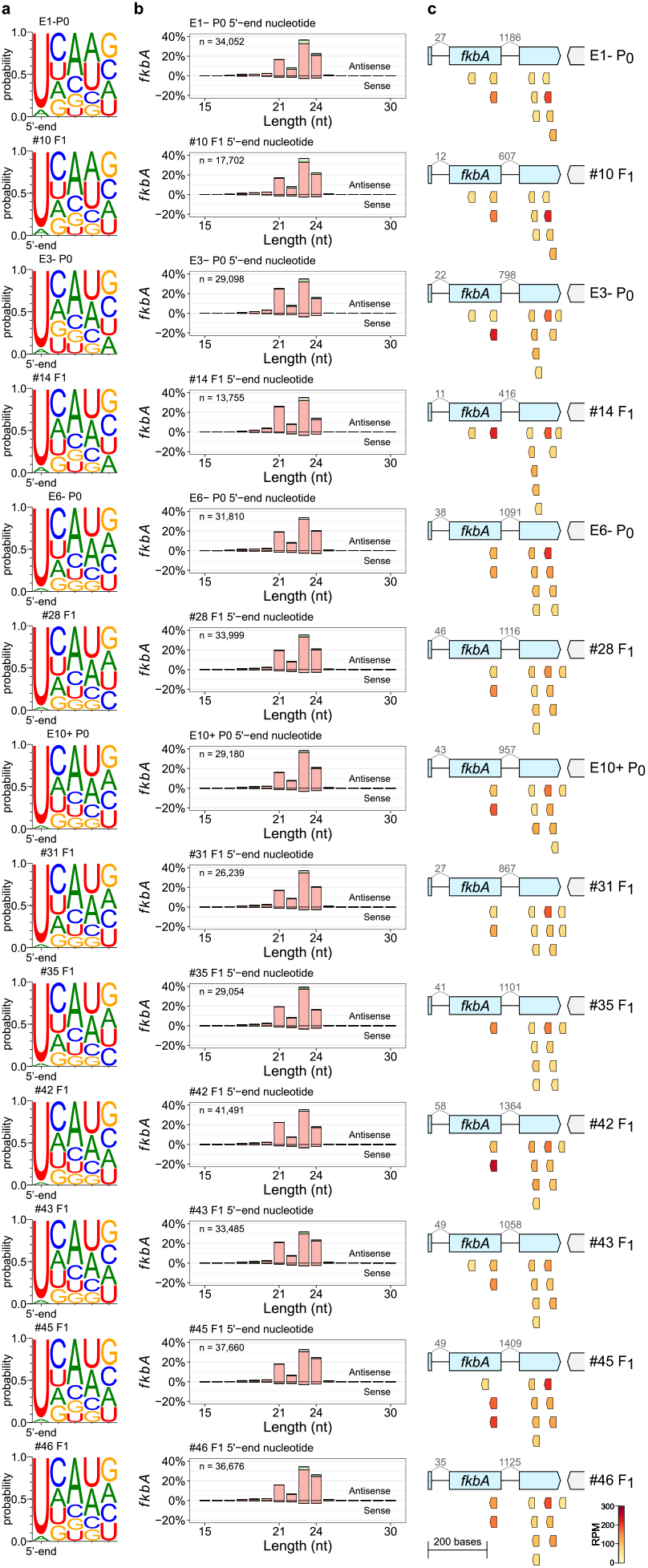
Inheritable siRNAs harbor common features and the most abundant molecules can be traced from progeny to their parents. **(a)** Sequence logos display the proportion of each nucleotide (color-coded) at the first five (5’-end) positions of the siRNA sequences found in the depicted epimutant parents and progeny. **(b)** The percentage of siRNA reads is plotted according to length, strand sense, and 5’-end nucleotide distribution. **(c)** The ten most abundant siRNA molecules in each epimutant are color-coded according to their relative abundance as reads per million (RPM) and plotted across the *fkbA* locus. In the gene model, the number of siRNA reads spanning across each exon-exon junction are displayed above curved lines (in grey) for each epimutant. Inheritable siRNA molecules exhibit lengths ranging from 21 to 24 nt, harbor a 5’-uracil, are antisense, and derive mainly from mature mRNAs. There is a notable correlation between the most abundant siRNAs found in parents and their progeny.

**Extended Data Fig. 10.**
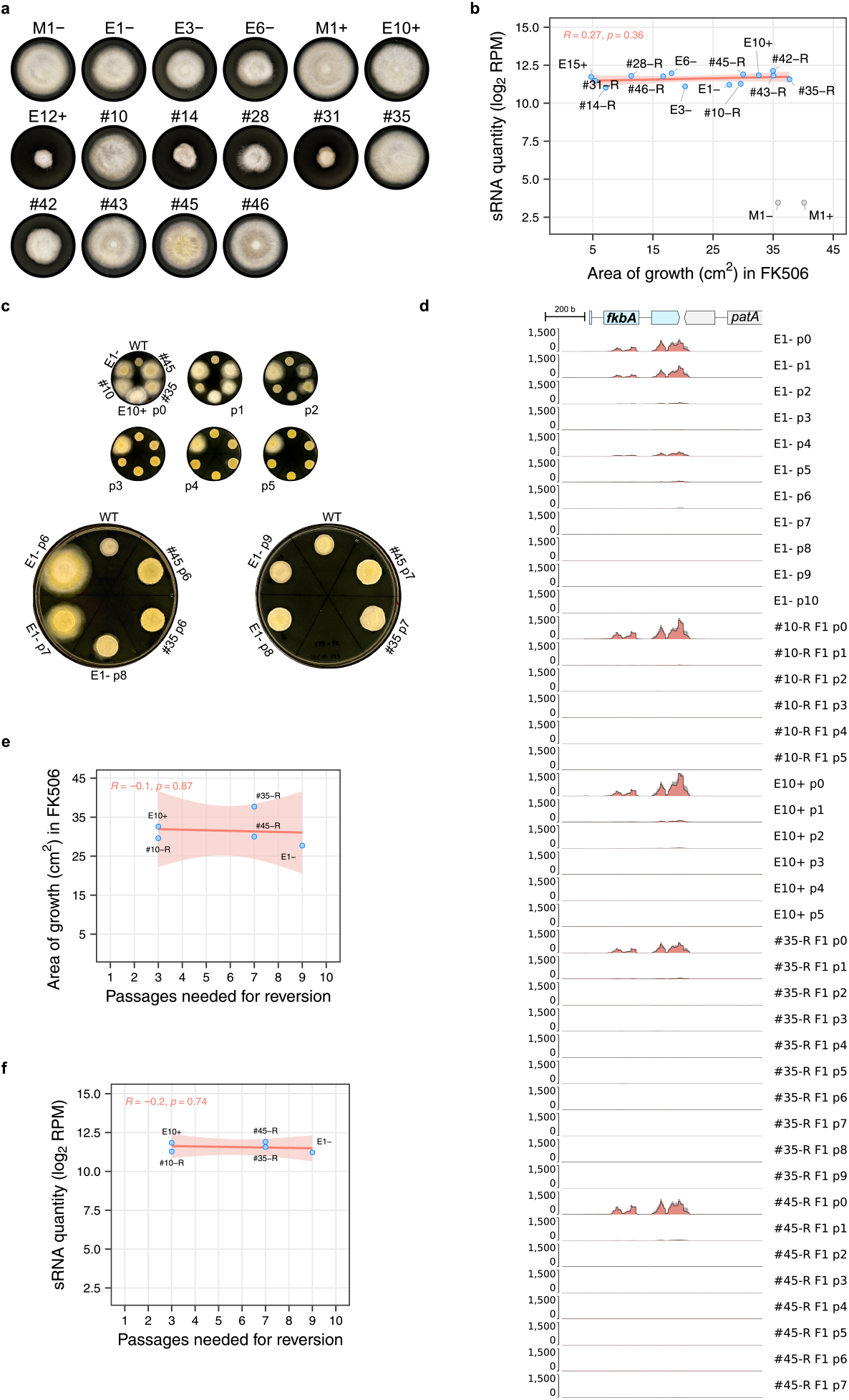
Progressive loss of FK506 and rapamycin resistance coincides with siRNA depletion. **(a)** Growth of representative resistant strains during 72-hour exposure to FK506 and rapamycin. **(b)** Dot plot showing the correlation between growth area (cm^2^), used as a proxy for the extent of resistance, and sRNA quantity, measured as normalized log_2_ reads per million (RPM). A red regression line with a shaded confidence interval is shown. Pearson’s R and the associated p-value are indicated in red. Although resistance levels vary among isolates, they do not correlate with sRNA quantity. **(c)** Growth of selected strains during FK506 exposure across successive drug-free vegetative passages until reversion to drug susceptibility. Passage number increases from left to right. **(d)** Genomic plot showing sRNA coverage (dark gray for total sRNA, red overlay for siRNA) across the *fkbA* locus in the corresponding drug-free passages shown in **(c). (e, f)** Dot plots showing correlations between the number of passages required for reversion and the extent of resistance (**e**) or sRNA quantity (**f**) in strains shown in **(d)**. A red regression line with a shaded confidence interval is shown. Pearson’s R and the p-value are indicated in red.

## References

1. E. M. Darby, E. Trampari, P. Siasat, M. S. Gaya, I. Alav, M. A. Webber, and J. M. A. Blair. Molecular mechanisms of antibiotic resistance revisited. Nature Reviews Microbiology, 21 (5):280–295, 2023. doi: 10.1038/s41579-022-00820-y.

2. Geneva: World Health Organization. WHO fungal priority pathogens list to guide research, development and public health action, 2022.

3. M. C. Fisher, F. Burnett, C. Chandler, N. A. R. Gow, S. Gurr, A. Hart, A. Holmes, R. C. May, J. Quinn, T. Soliman, N. J. Talbot, H. M. West, J. S. West, P. L. White, M. Bromley, and D. Armstrong-James. A one health roadmap towards understanding and mitigating emerging Fungal Antimicrobial Resistance: fAMR. npj Antimicrobials and Resistance, 2(1): 36, 2024. doi: 10.1038/s44259-024-00055-2.

4. M. C. Fisher, A. Alastruey-Izquierdo, J. Berman, T. Bicanic, E. M. Bignell, P. Bowyer, M. Bromley, R. Brüggemann, G. Garber, O. A. Cornely, S. J. Gurr, T. S. Harrison, E. Kuijper, J. Rhodes, D. C. Sheppard, A. Warris, P. L. White, J. Xu, B. Zwaan, and P. E. Verweij. Tackling the emerging threat of antifungal resistance to human health. Nature Reviews Microbiology, 20(9):557–571, 2022. doi: 10.1038/s41579-022-00720-1.

5. N. A. R. Gow, C. Johnson, J. Berman, A. T. Coste, C. A. Cuomo, D. S. Perlin, T. Bicanic, T. S. Harrison, N. Wiederhold, M. Bromley, T. Chiller, and K. Edgar. The importance of antimicrobial resistance in medical mycology. Nature Communications, 13(1):5352, 2022. doi: 10.1038/s41467-022-32249-5..:

6. S. Calo, C. Shertz-Wall, S. Lee, R. Bastidas, F. Nicolás, J. Granek, P. Mieczkowski, S. Torres-Martínez, R. Ruiz-Vázquez, M. Cardenas, and J. Heitman. Antifungal drug resistance evoked via RNAi-dependent epimutations. Nature, 513:555–558, 2014. doi: 10.1038/nature13575..:

7. Z. Chang and J. Heitman. Drug-resistant epimutants exhibit organ-specific stability and induction during murine infections caused by the human fungal pathogen Mucor circinelloides. mBio, 10(6):02579–19, 2019. doi: 10.1128/mBio.02579-19..:

8. S. Torres-Garcia, I. Yaseen, M. Shukla, P. Audergon, S. White, A. Pidoux, and R. Allshire. Epigenetic gene silencing by heterochromatin primes fungal resistance. Nature, 585(7825): 453–458, 2020. doi: 10.1038/s41586-020-2706-x..:

9. H. Kramer, D. Cook, M. Seidl, and B. Thomma. Epigenetic regulation of nuclear processes in fungal plant pathogens. PLOS Pathogens, 19(8):1011525, 2023. doi: 10.1371/journal.ppat.1011525..:

10. E. Heard and R. Martienssen. Transgenerational epigenetic inheritance: myths and mechanisms. Cell, 157(1):95–109, 2014. doi: 10.1016/j.cell.2014.02.045..:

11. K. Skvortsova, N. Iovino, and O. Bogdanović. Functions and mechanisms of epigenetic inheritance in animals. Nature Reviews Molecular Cell Biology, 19(12):774–790, 2018. doi: 10.1038/s41580-018-0074-2..:

12. M. Fitz-James and G. Cavalli. Molecular mechanisms of transgenerational epigenetic inheritance. Nature Reviews Genetics, 23(6):325–341, 2022. doi: 10.1038/s41576-021-00438-5..:

13. L. Wagner, J. Stielow, G. Hoog, K. Bensch, V. Schwartze, K. Voigt, A. Alastruey-Izquierdo, O. Kurzai, and G. Walther. A new species concept for the clinically relevant Mucor circinelloides complex. Persoonia, 44:67–97, 2020. doi: 10.3767/persoonia.2020.44.03..:

14. R.M. Ruiz-Vázquez, F. E. Nicolás, S. Torres-Martínez, and V. Garre. Distinct RNAi pathways in the regulation of physiology and development in the fungus Mucor circinelloides. In T. Friedmann, J. C. Dunlap, and S. F. Goodwin, editors, Advances in Genetics, volume 91, pages 55–102. Academic Press, 2015. ISBN 9780128029213. doi: 10.1016/bs.adgen.2015.07.002..:

15. S. Calo, F. Nicolás, S. Lee, A. Vila, M. Cervantes, S. Torres-Martínez, R. Ruiz-Vázquez,.: M. Cardenas, and J. Heitman. A non-canonical RNA degradation pathway suppresses RNAi-dependent epimutations in the human fungal pathogen Mucor circinelloides. PLoS Genetics, 13(3):1006686, 2017. doi: 10.1371/journal.pgen.1006686.

16. C. Pérez-Arques, M. Navarro-Mendoza, L. Murcia, E. Navarro, V. Garre, and F. Nicolás. A non-canonical RNAi pathway controls virulence and genome stability in Mucorales. PLoS Genetics, 16(7):1008611, 2020. doi: 10.1371/journal.pgen.1008611..:

17. R. J. Bastidas, C. A. Shertz, S. C. Lee, J. Heitman, and M. E. Cardenas. Rapamycin exerts antifungal activity in vitro and in vivo against <i>Mucor circinelloides</i> via FKBP12-dependent inhibition of TOR. Eukaryotic cell, 11(3):270–281, 2012. doi: 10.1128/EC. 05284–11..:

18. Z. Chang, R. Billmyre, S. Lee, and J. Heitman. Broad antifungal resistance mediated by RNAi-dependent epimutation in the basal human fungal pathogen Mucor circinelloides. PLoS Genetics, 15(2):1007957, 2019. doi: 10.1371/journal.pgen.1007957..:

19. A. Blakeslee. Sexual reproduction in the Mucorineae. Proceedings of the American Academy of Arts and Sciences, 40(4):205–319, 1904. doi: 10.2307/20021962..:

20. A. Eslava, M. Alvarez, and M. Delbrück. Meiosis in Phycomyces. Proceedings of the National Academy of Sciences, 72(10):4076–4080, 1975. doi: 10.1073/pnas.72.10.4076..:

21. E. Cerdá-Olmedo. The genetics of Phycomyces blakesleeanus. Genetics Research, 25(3): 285–296, 1975. doi: 10.1017/S0016672300015718..:

22. W. Gauger. The germination of zygospores of Mucor hiemalis. Mycologia, 57(4):634–641, 1965. doi: 10.2307/3756739..:

23. F. R. James and W. Gauger. Studies on the genetics of Mucor hiemalis. Mycologia, 74(5): 744, 1982. doi: 10.2307/3792860..:

24. D. Qutob, B. Patrick Chapman, and M. Gijzen. Transgenerational gene silencing causes gain of virulence in a plant pathogen. Nature Communications, 4(1):1349, 2013. doi: 10.1038/ncomms2354..:

25. S. Mondo, R. Dannebaum, R. Kuo, K. Louie, A. Bewick, K. LaButti, S. Haridas, A. Kuo, Salamov, S. Ahrendt, R. Lau, B. Bowen, A. Lipzen, W. Sullivan, B. Andreopoulos, A. Clum, E. Lindquist, C. Daum, T. Northen, and I. Grigoriev. Widespread adenine N6-methylation of active genes in fungi. Nature Genetics, 49(6):964–968, 2017. doi: 10.1038/ng.3859.

26. M. Navarro-Mendoza, C. Pérez-Arques, and J. Heitman. Heterochromatin and RNAi act independently to ensure genome stability in Mucorales human fungal pathogens. Proceedings of the National Academy of Sciences, 120(7):2220475120, 2023. doi: 10.1073/pnas.2220475120..:

27. C. Lax, S. Mondo, M. Osorio-Concepción, A. Muszewska, M. Corrochano-Luque, G. Gutiérrez, R. Riley, A. Lipzen, J. Guo, H. Hundley, M. Amirebrahimi, V. Ng, D. Lorenzo-Gutiérrez,.: U. Binder, J. Yang, Y. Song, D. Cánovas, E. Navarro, M. Freitag, and V. Garre. Symmetric and asymmetric DNA N6-adenine methylation regulates different biological responses in Mucorales. Nature Communications, 15(1):6066, 2024. doi: 10.1038/s41467-024-50365-2.

28. P. Higgs and N. Lehman. The RNA World: molecular cooperation at the origins of life. Nature Reviews Genetics, 16(1):7–17, 2015. doi: 10.1038/nrg3841..:

29. E. Koonin and V. Dolja. Virus world as an evolutionary network of viruses and capsidless selfish elements. Microbiology and Molecular Biology Reviews, 78(2):278–303, 2014. doi: 10.1128/MMBR.00049-13..:

30. L. Li, P. Zheng, and J. Dean. Maternal control of early mouse development. Development, 137(6):859–870, 2010. doi: 10.1242/dev.039487..:

31. K. Gapp, A. Jawaid, P. Sarkies, J. Bohacek, P. Pelczar, J. Prados, L. Farinelli, E. Miska, and I. Mansuy. Implication of sperm RNAs in transgenerational inheritance of the effects of early trauma in mice. Nature Neuroscience, 17(5):667–669, 2014. doi: 10.1038/nn.3695.

32. Q. Chen, M. Yan, Z. Cao, X. Li, Y. Zhang, J. Shi, G. Feng, H. Peng, X. Zhang, Y. Zhang, J. Qian, E. Duan, Q. Zhai, and Q. Zhou. Sperm tsRNAs contribute to intergenerational inheritance of an acquired metabolic disorder. Science, 351(6271):397–400, 2016. doi: 10.1126/science.aad7977.

33. I. Rieger, G. Weintraub, I. Lev, K. Goldstein, D. Bar-Zvi, S. Anava, H. Gingold, S. Shaham, and O. Rechavi. Nucleus-independent transgenerational small RNA inheritance in Caenorhabditis elegans. Science Advances, 9(43), 2023. doi: 10.1126/sciadv.adj8618..:

34. R. Wilson, M. Le Bourgeois, M. Perez, and P. Sarkies. Fluctuations in chromatin state at regulatory loci occur spontaneously under relaxed selection and are associated with epigenetically inherited variation in C. elegans gene expression. PLoS Genetics, 19(3):1010647, 2023. doi: 10.1371/journal.pgen.1010647..:

35. S. Grewal. The molecular basis of heterochromatin assembly and epigenetic inheritance.Molecular Cell, 83(11):1767–1785, 2023. doi: 10.1016/j.molcel.2023.04.020.

36. S. I. Grewal and A. J. Klar. Chromosomal inheritance of epigenetic states in fission.: east during mitosis and meiosis. Cell, 86(1):95–101, 1996. doi: 10.1016/S0092-8674(00)80080-X.

37. P. N. C. B. Audergon, S. Catania, A. Kagansky, P. Tong, M. Shukla, A. L. Pidoux, and R. C. Allshire. Restricted epigenetic inheritance of H3K9 methylation. Science, 348(6230):132–135, 2015. doi: 10.1126/science.1260638..:

38. K. Ragunathan, G. Jih, and D. Moazed. Epigenetic inheritance uncoupled from sequence-specific recruitment. Science, 348(6230), 2015. doi: 10.1126/science.1258699.

39. E. Schulz, J. Wetzel, A. Burmester, S. Ellenberger, L. Siegmund, and J. Wöstemeyer. Sex loci of homothallic and heterothallic Mucorales. Endocytobiosis and Cell Research, 27(4): 39–57, 2016.

40. S. Calo, F. Nicolás, A. Vila, S. Torres-Martínez, and R. Ruiz-Vázquez. Two distinct RNA-dependent RNA polymerases are required for initiation and amplification of RNA silencing in the basal fungus Mucor circinelloides. Molecular Microbiology, 83(2):379–394, 2012. doi: 10.1111/j.1365-2958.2011.07939.x.

41. F. Nicolás, A. Vila, S. Moxon, M. Cascales, S. Torres-Martínez, R. Ruiz-Vázquez, and V. Garre. The RNAi machinery controls distinct responses to environmental signals in the basal fungus Mucor circinelloides. BMC Genomics, 16(1):237, 2015. doi: 10.1186/s12864-015-1443-2.

42. T. Beltran, V. Shahrezaei, V. Katju, and P. Sarkies. Epimutations driven by small RNAs arise frequently but most have limited duration in Caenorhabditis elegans. Nature Ecology & Evolution, 4(11):1539–1548, 2020. doi: 10.1038/s41559-020-01293-z.

43. A. Grishok, H. Tabara, and C. Mello. Genetic requirements for inheritance of RNAi in C. elegans. Science, 287(5462):2494–2497, 2000. doi: 10.1126/science.287.5462.2494.

44. M. Duraisingh and K. Skillman. Epigenetic variation and regulation in malaria parasites. Annual Review of Microbiology, 72(1):355–375, 2018. doi: 10.1146/annurev-micro-090817-062722.

45. A. Buscaino. Chromatin-mediated regulation of genome plasticity in human fungal pathogens. Genes, 10(11):855, 2019. doi: 10.3390/genes10110855.

46. F. Nicolás, S. Torres-Martínez, and R. Ruiz-Vázquez. Loss and retention of RNA interference in fungi and parasites. PLoS Pathogens, 9(1):1003089, 2013. doi: 10.1371/journal.ppat.1003089.

47. E. Iracane, C. Arias-Sardá, C. Maufrais, I. Ene, C. D’Enfert, and A. Buscaino. Identification of an active RNAi pathway in Candida albicans. Proceedings of the National Academy of Sciences, 121(17), 2024. doi: 10.1073/pnas.2315926121.

48. C. Pérez-Arques, M. Navarro-Mendoza, L. Murcia, E. Navarro, V. Garre, and F. Nicolás. The RNAi mechanism regulates a new exonuclease gene involved in the virulence of Mucorales. International Journal of Molecular Sciences, 22(5):2282, 2021. doi: 10.3390/ijms22052282.

49. M. Navarro-Mendoza, C. Pérez-Arques, S. Panchal, F. Nicolás, S. Mondo, P. Ganguly, J. Pangilinan, I. Grigoriev, J. Heitman, K. Sanyal, and V. Garre. Early diverging fungus Mucor circinelloides lacks centromeric histone CENP-A and displays a mosaic of point and regional centromeres. Current Biology, 29(22):3791–3802, 2019. doi: 10.1016/j.cub.2019.09.024.

50. M. Kolmogorov, J. Yuan, Y. Lin, and P. Pevzner. Assembly of long, error-prone reads using repeat graphs. Nature Biotechnology, 37(5):540–546, 2019. doi: 10.1038/s41587-019-0072-8.

51. R. Vaser, I. Sović, N. Nagarajan, and M. Šikić. Fast and accurate de novo genome assembly from long uncorrected reads. Genome Research, 27(5):737–746, 2017. doi: 10.1101/gr.214270.116.

52. J. Hu, J. Fan, Z. Sun, and S. Liu. NextPolish: a fast and efficient genome polishing tool for long-read assembly. Bioinformatics, 36(7):2253–2255, 2020. doi: 10.1093/bioinformatics/btz891.

53. J. Hu, Z. Wang, F. Liang, S.-L. Liu, K. Ye, and D.-P. Wang. NextPolish2: A repeat-aware polishing tool for genomes assembled using HiFi long reads. Genomics, Proteomics & Bioinformatics, 22(1), 2024. doi: 10.1093/gpbjnl/qzad009.

54. H. Li. Minimap2: pairwise alignment for nucleotide sequences. Bioinformatics, 34(18): 3094–3100, 2018. doi: 10.1093/bioinformatics/bty191.

55. M. Krzywinski, J. Schein, I. Birol, J. Connors, R. Gascoyne, D. Horsman, S. J. Jones, and M. A. Marra. Circos: an information aesthetic for comparative genomics. Genome research, 19(9):1639–1645, 2009. doi: 10.1101/gr.092759.109.

56. J. Flynn, R. Hubley, C. Goubert, J. Rosen, A. Clark, C. Feschotte, and A. Smit. RepeatModeler2 for automated genomic discovery of transposable element families. Proceedings of the National Academy of Sciences, 117(17):9451–9457, 2020. doi: 10.1073/pnas.1921046117.

57. W. Bao, K. Kojima, and O. Kohany. Repbase Update, a database of repetitive elements in eukaryotic genomes. Mobile DNA, 6(1):11, 2015. doi: 10.1186/s13100-015-0041-9.

58. J. Palmer and J. Stajich. Funannotate v1.8.1: Eukaryotic genome annotation, 2020. Zenodo.

59. A. Gurevich, V. Saveliev, N. Vyahhi, and G. Tesler. QUAST: quality assessment tool for genome assemblies. Bioinformatics, 29(8):1072–1075, 2013. doi: 10.1093/bioinformatics/btt086.

60. M. Manni, M. Berkeley, M. Seppey, F. Simão, and E. Zdobnov. BUSCO update: novel and streamlined workflows along with broader and deeper phylogenetic coverage for scoring of eukaryotic, prokaryotic, and viral genomes. Molecular Biology and Evolution, 38(10): 4647–4654, 2021. doi: 10.1093/molbev/msab199.

61. C. Camacho, G. Coulouris, V. Avagyan, N. Ma, J. Papadopoulos, K. Bealer, and T. Madden. BLAST+: architecture and applications. BMC Bioinformatics, 10(1):421, 2009. doi: 10.1186/1471-2105-10-421.

62. P. Jones, D. Binns, H.-Y. Chang, M. Fraser, W. Li, C. McAnulla, H. McWilliam, J. Maslen, Mitchell, G. Nuka, S. Pesseat, A. Quinn, A. Sangrador-Vegas, M. Scheremetjew, S.-Y. Yong, R. Lopez, and S. Hunter. InterProScan 5: genome-scale protein function classification. Bioinformatics, 30(9):1236–1240, 2014. doi: 10.1093/bioinformatics/btu031.

63. S. Chen, Y. Zhou, Y. Chen, and J. Gu. fastp: an ultra-fast all-in-one FASTQ preprocessor. Bioinformatics, 34(17):884–890, 2018. doi: 10.1093/bioinformatics/bty560.

64. H. Li and R. Durbin. Fast and accurate short read alignment with Burrows-Wheeler transform. Bioinformatics, 25(14):1754–1760, 2009. doi: 10.1093/bioinformatics/btp324.

65. G. Auwera and B. O’Connor. Genomics in the Cloud: Using Docker, GATK, and WDL in Terra. O’Reilly Media, Inc, 2020.

66. A. Quinlan and I. Hall. BEDTools: a flexible suite of utilities for comparing genomic features. Bioinformatics, 26(6):841–842, 2010. doi: 10.1093/bioinformatics/btq033.

67. F. Ramírez, D. Ryan, B. Grüning, V. Bhardwaj, F. Kilpert, A. Richter, S. Heyne, F. Dündar, and.: T. Manke. deepTools2: a next generation web server for deep-sequencing data analysis. Nucleic Acids Research, 44(W1):160–165, 2016. doi: 10.1093/nar/gkw257.

68. B. Langmead, C. Trapnell, M. Pop, and S. Salzberg. Ultrafast and memory-efficient alignment of short DNA sequences to the human genome. Genome Biology, 10(3):25, 2009. doi: 10.1186/gb-2009-10-3-r25.

69. A. Dobin and T. Gingeras. Mapping RNA-seq Reads with STAR. In Current Protocols in Bioinformatics, page 11 14 1–11 14 19. John Wiley & Sons, Inc, 2015. doi: 10.1002/0471250953.bi1114s51.

70. H. Li, B. Handsaker, A. Wysoker, T. Fennell, J. Ruan, N. Homer, G. Marth, G. Abecasis, and R. Durbin. The Sequence Alignment/Map format and SAMtools. Bioinformatics, 25(16): 2078–2079, 2009. doi: 10.1093/bioinformatics/btp352.

71. M. I. Love, W. Huber, and S. Anders. Moderated estimation of fold change and dispersion for RNA-seq data with DESeq2. Genome Biology, 15(12):550, 2014. doi: 10.1186/s13059-014-0550-8.

72. Z. Gu. Complex heatmap visualization. iMeta, 1(3), 2022. doi: 10.1002/imt2.43.

73. Y. Zhang, T. Liu, C. Meyer, J. Eeckhoute, D. Johnson, B. Bernstein, C. Nussbaum, R. Myers, M. Brown, W. Li, and X. Liu. Model-based Analysis of ChIP-Seq (MACS). Genome Biology, 9(9):137, 2008. doi: 10.1186/gb-2008-9-9-r137.

74. C. S. Ross-Innes, R. Stark, A. E. Teschendorff, K. A. Holmes, H. R. Ali, M. J. Dunning, G. D. Brown, O. Gojis, I. O. Ellis, A. R. Green, S. Ali, S.-F. Chin, C. Palmieri, C. Caldas, and J. S. Carroll. Differential oestrogen receptor binding is associated with clinical outcome in breast cancer. Nature, 481(7381):389–393, 2012. doi: 10.1038/nature10730.

75. C. Gao, C. Chen, T. Akyol, A. Dusa, G. Yu, B. Cao, and P. Cai. ggVennDiagram: Intuitive Venn diagram software extended. iMeta, 3(1), 2024. doi: 10.1002/imt2.177.

76. T. Hothorn, F. Bretz, and P. Westfall. Simultaneous inference in general parametric models. Biometrical Journal, 50(3):346–363, 2008. doi: 10.1002/bimj.200810425.

