## Supplementary information for "RNAi epimutations conferring antifungal drug resistance are inheritable"

#### Extended Data Table 1

| Assembly metrics | *M. circinelloides* PS15− | *M. circinelloides* PS15+ | *M. lusitanicus* CBS277.49 v3.0 |
|---|---|---|---|
| **Total length** | 37,441,900 | 37,217,483 | 36,611,222 |
| **# contigs** | 14 | 15 | 20 (12 scaffolds) |
| **Percent gaps** | 0.000% | 0.000% | 0.011% |
| **Largest contig** | 4,896,228 | 4,875,796 | 6,403,782 |
| **N50** | 3,291,799 | 2,902,562 | 4,607,068 |
| **N90** | 1,855,953 | 1,865,875 | 2,643,553 |
| **L50** | 5 | 5 | 4 |
| **L90** | 11 | 11 | 8 |
| **GC content** | 39.42% | 39.41% | 42.18% |
| **# genes** | 12,397 | 12,325 | 11,616 |
| **BUSCO completeness** | 97.3% | 97.3% | 94.9% |

**Extended Data Table 1.** Genome assembly metrics for *M. circinelloides* PS15 opposite mating types. Values are compared to the latest *M. lusitanicus* CBS277.49 genome assembly. Assembly completeness is assessed as the percentage of conserved orthologs from the BUSCO Eukaryota ortholog database (eukaryota_odb10).

#### Extended Data Table 2

| Isolate | Derived from | Phenotype | Cause | Reference |
|---|---|---|---|---|
| PS15– | *M. circinelloides* CBS394.68 | FK506/rapamycin susceptible | | (4) |
| PS15+ | *M. circinelloides* CBS172.27 | FK506/rapamycin susceptible | | (4) |
| M1- | PS15- | FK506/rapamycin resistant | *fkbA* c.112delC, p.(Lys17ArgfsTer58) | This study |
| E1- | PS15- | FK506/rapamycin resistant | *fkbA* epimutation | This study |
| E2- | PS15- | FK506/rapamycin resistant | *fkbA* epimutation | This study |
| E3- | PS15- | FK506/rapamycin resistant | *fkbA* epimutation | This study |
| E4- | PS15- | FK506/rapamycin resistant | *fkbA* epimutation | This study |
| E5- | PS15- | FK506/rapamycin resistant | *fkbA* epimutation | This study |
| E6- | PS15- | FK506/rapamycin resistant | *fkbA* epimutation | This study |
| M1+ | PS15+ | FK506/rapamycin resistant | *fkbA* c.119_120delGG, p.(Gly20ArgfsTer15) | This study |
| M2+ | PS15+ | FK506/rapamycin resistant | *fkbA* c.205delA, p.(Gln48HisfsTer28) | This study |
| M3+ | PS15+ | FK506/rapamycin resistant | *fkbA* c.112delC, p.(Lys17ArgfsTer58) | This study |
| E10+ | PS15+ | FK506/rapamycin resistant | *fkbA* epimutation | This study |
| E12+ | PS15+ | FK506/rapamycin resistant | *fkbA* epimutation | This study |
| E15+ | PS15+ | FK506/rapamycin resistant | *fkbA* epimutation | This study |
| #1 | PS15– and PS15+ cross | FK506/rapamycin susceptible | | This study |
| #2 | PS15– and PS15+ cross | FK506/rapamycin susceptible | | This study |
| #3 | PS15– and PS15+ cross | FK506/rapamycin susceptible | | This study |
| #4 | PS15– and PS15+ cross | FK506/rapamycin susceptible | | This study |
| #5 | PS15– and PS15+ cross | FK506/rapamycin susceptible | | This study |
| #6 | PS15– and PS15+ cross | FK506/rapamycin susceptible | | This study |
| #7 | PS15– and PS15+ cross | FK506/rapamycin susceptible | | This study |
| #8 | PS15– and PS15+ cross | FK506/rapamycin susceptible | | This study |
| #9 | E1– and PS15+ cross | FK506/rapamycin susceptible | | This study |
| #10 | E1– and PS15+ cross | FK506/rapamycin resistant | *fkbA* epimutation | This study |
| #11 | E1– and PS15+ cross | FK506/rapamycin susceptible | | This study |
| #12 | E1– and PS15+ cross | FK506/rapamycin susceptible | | This study |
| #13 | E3– and PS15+ cross | FK506/rapamycin susceptible | | This study |
| #14 | E3– and PS15+ cross | FK506/rapamycin resistant | *fkbA* epimutation | This study |
| #15 | E3– and PS15+ cross | FK506/rapamycin susceptible | | This study |
| #16 | E3– and PS15+ cross | FK506/rapamycin susceptible | | This study |
| #17 | E3– and PS15+ cross | FK506/rapamycin susceptible | | This study |
| #18 | E3– and PS15+ cross | FK506/rapamycin susceptible | | This study |
| #19 | E3– and PS15+ cross | FK506/rapamycin susceptible | | This study |
| #20 | E3– and PS15+ cross | FK506/rapamycin susceptible | | This study |
| #21 | E3– and PS15+ cross | FK506/rapamycin susceptible | | This study |
| #22 | E4– and PS15+ cross | FK506/rapamycin susceptible | | This study |
| #23 | E4– and PS15+ cross | FK506/rapamycin susceptible | | This study |
| #24 | E4– and PS15+ cross | FK506/rapamycin resistant | *fkbA* epimutation | This study |
| #25 | E4– and PS15+ cross | FK506/rapamycin susceptible | | This study |
| #26 | E6– and PS15+ cross | FK506/rapamycin resistant | *fkbA* epimutation | This study |
| #27 | E6– and PS15+ cross | FK506/rapamycin susceptible | | This study |
| #28 | E6– and PS15+ cross | FK506/rapamycin resistant | *fkbA* epimutation | This study |
| #29 | E6– and PS15+ cross | FK506/rapamycin susceptible | | This study |
| #30 | E6– and PS15+ cross | FK506/rapamycin resistant | *fkbA* epimutation | This study |
| #31 | PS15– and E10+ cross | FK506/rapamycin resistant | *fkbA* epimutation | This study |
| #32 | PS15– and E10+ cross | FK506/rapamycin susceptible | | This study |
| #33 | PS15– and E10+ cross | FK506/rapamycin susceptible | | This study |
| #34 | PS15– and E10+ cross | FK506/rapamycin susceptible | | This study |
| #35 | PS15– and E10+ cross | FK506/rapamycin resistant | *fkbA* epimutation | This study |
| #36 | PS15– and E10+ cross | FK506/rapamycin susceptible | | This study |
| #37 | PS15– and E10+ cross | FK506/rapamycin susceptible | | This study |
| #38 | PS15– and E10+ cross | FK506/rapamycin susceptible | | This study |
| #39 | PS15– and E15+ cross | FK506/rapamycin susceptible | | This study |
| #40 | PS15– and E15+ cross | FK506/rapamycin susceptible | | This study |
| #41 | PS15– and E15+ cross | FK506/rapamycin susceptible | | This study |
| #42 | E1– and E10+ cross | FK506/rapamycin resistant | *fkbA* epimutation | This study |
| #43 | E1– and E10+ cross | FK506/rapamycin resistant | *fkbA* epimutation | This study |
| #44 | E1– and E10+ cross | FK506/rapamycin susceptible | | This study |
| #45 | E1– and E10+ cross | FK506/rapamycin resistant | *fkbA* epimutation | This study |
| #46 | E1– and E10+ cross | FK506/rapamycin resistant | *fkbA* epimutation | This study |
| #47 | E3– and E15+ cross | FK506/rapamycin susceptible | | This study |
| #48 | E3– and E15+ cross | FK506/rapamycin resistant | *fkbA* epimutation | This study |
| #49 | E3– and E15+ cross | FK506/rapamycin resistant | *fkbA* epimutation | This study |
| #50 | E3– and E15+ cross | FK506/rapamycin resistant | *fkbA* epimutation | This study |
| #51 | PS15– and M1+ cross | FK506/rapamycin resistant | M1+ *fkbA* mutation | This study |
| #52 | PS15– and M1+ cross | FK506/rapamycin resistant | M1+ *fkbA* mutation | This study |
| #53 | PS15– and M1+ cross | FK506/rapamycin resistant | M1+ *fkbA* mutation | This study |
| #54 | PS15– and M1+ cross | FK506/rapamycin susceptible | | This study |
| #55 | PS15– and M1+ cross | FK506/rapamycin susceptible | | This study |
| PS10 WT | *M. lusitanicus* R7B | FK506/rapamycin susceptible | | (1) |
| PS10 E1 | *M. lusitanicus* R7B | FK506/rapamycin resistant | *fkbA* epimutation | (1) |
| PS10 WT | *M. lusitanicus* MU402 *rdrp1*Δ | 5-FOA susceptible, Rdrp1- | | (17) |
| PS10 E3 | *M. lusitanicus* MU402 *rdrp1*Δ | 5-FOA resistant, , Rdrp1- | *pyrF* epimutation | (6) |
| PS10 E4 | *M. lusitanicus* MU402 *rdrp1*Δ | 5-FOA resistant, Rdrp1- | *pyrG* epimutation | (6) |
| PS14 WT | *M. circinelloides* 1006PhL | FK506/rapamycin susceptible | | (1) |
| PS14 E3 | *M. circinelloides* 1006PhL | FK506/rapamycin resistant | *fkbA* epimutation | (1) |

**Extended Data Table S2.** Strains utilized in this work are listed highlighting the isolate name in the manuscript, their parental strains, relevant phenotype, cause of drug resistance, and reference.

#### Extended Data Table 3

| Name | Sequence (5'⟶3') | Description |
|---|---|---|
| JOHE51779 | AAAACGCGAAACAGCAAACG | *fkbA* whole-sequence amplification, Sanger-sequencing |
| JOHE51780 | GGTGACTTTGGTGGTTTACG | *fkbA* whole-sequence amplification, Sanger-sequencing |
| JOHE51662 | GGTGACAAAGTTACCATCCAC | *fkbA* Sanger-sequencing |
| JOHE51655 | CCTTGATGACTTGACCAACACC | *fkbA* Sanger-sequencing |
| JOHE51656 | taatacgactcactatagggatgggtgttacTGTTGAAAGAATTGCTCCT | *fkbA* template for in vitro transcription (small RNA probe) |
| JOHE51664 | AGGAATGAGGCCGGGGTAAC | *fkbA* template for in vitro transcription (small RNA probe) |
| JOHE51679 | CAACTTTTAACAACGGATCTC | *rRNA 5.8S* template for in vitro transcription (small RNA probe) |
| JOHE51680 | taatacgactcactataggggCTGAAACAGGCGTGCTCATTG | *rRNA 5.8S* template for in vitro transcription (small RNA probe) |
| JOHE51775 | TCCGTGACTTTAGTTCCAAGC | *pyrG* 5'-UTR amplicon for qPCR |
| JOHE51776 | TCCACGTTCGCTATAAGTCTTG | *pyrG* 5'-UTR amplicon for qPCR |
| JOHE51777 | CGTTGTCATGTCCCCTGGTG | *pyrG* exon amplicon for qPCR |
| JOHE51778 | GCCGACAATGATGATATCTCCTC | *pyrG* exon amplicon for qPCR |
| JOHE51773 | AAAGAGTGCGTCTCCACAAG | *pyrG* 3'-UTR amplicon for qPCR |
| JOHE51774 | TCTCTTATGTGGCTGATTGGAC | *pyrG* 3'-UTR amplicon for qPCR |
| JOHE51769/CPA139 | TCCACATGCCCTTTATAAACTTG | *pyrF* 5'-UTR amplicon for qPCR |
| JOHE51770/CPA140 | ACGTTGATAAGCCTTCATTATGG | *pyrF* 5'-UTR amplicon for qPCR |
| JOHE51771/CPA141 | GCCGCACCTCACCTTACTTC | *pyrF* exon amplicon for qPCR |
| JOHE51772/CPA142 | TAGTCAAAGCCGGCATCCTC | *pyrF* exon amplicon for qPCR |
| JOHE51789/CPA159 | AGGATTAGAGCCGTCGTTATTG | *pyrF* 3'-UTR amplicon for qPCR |
| JOHE51768/CPA138 | AGATCCGGGTTTCACATGGC | *pyrF* 3'-UTR amplicon for qPCR |
| JOHE51718 | GTCTTGTCTCATGCAGACT | *GremLINE1* amplicon for qPCR, H3K9me2 positive control |
| JOHE51719 | GCACCAAGAGAGAGACGAGC | *GremLINE1* amplicon for qPCR, H3K9me2 positive control |
| JOHE51668 | ACCTTGGTGCTCGTCTTGCC | *vma1* amplicon for qPCR, H3K9me2 negative control normalization |
| JOHE51666 | CAGCACCAACAACAGTGACG | *vma1* amplicon for qPCR, H3K9me2 negative control normalization |

**Extended Data Table 3.** Primers used for this study are listed including their sequence, and brief description. Lowercase characters indicate primer tails containing the T7 promoter sequence (underlined) and a short spacer sequence for in vitro riboprobe transcription.

Extended Data Fig. 1

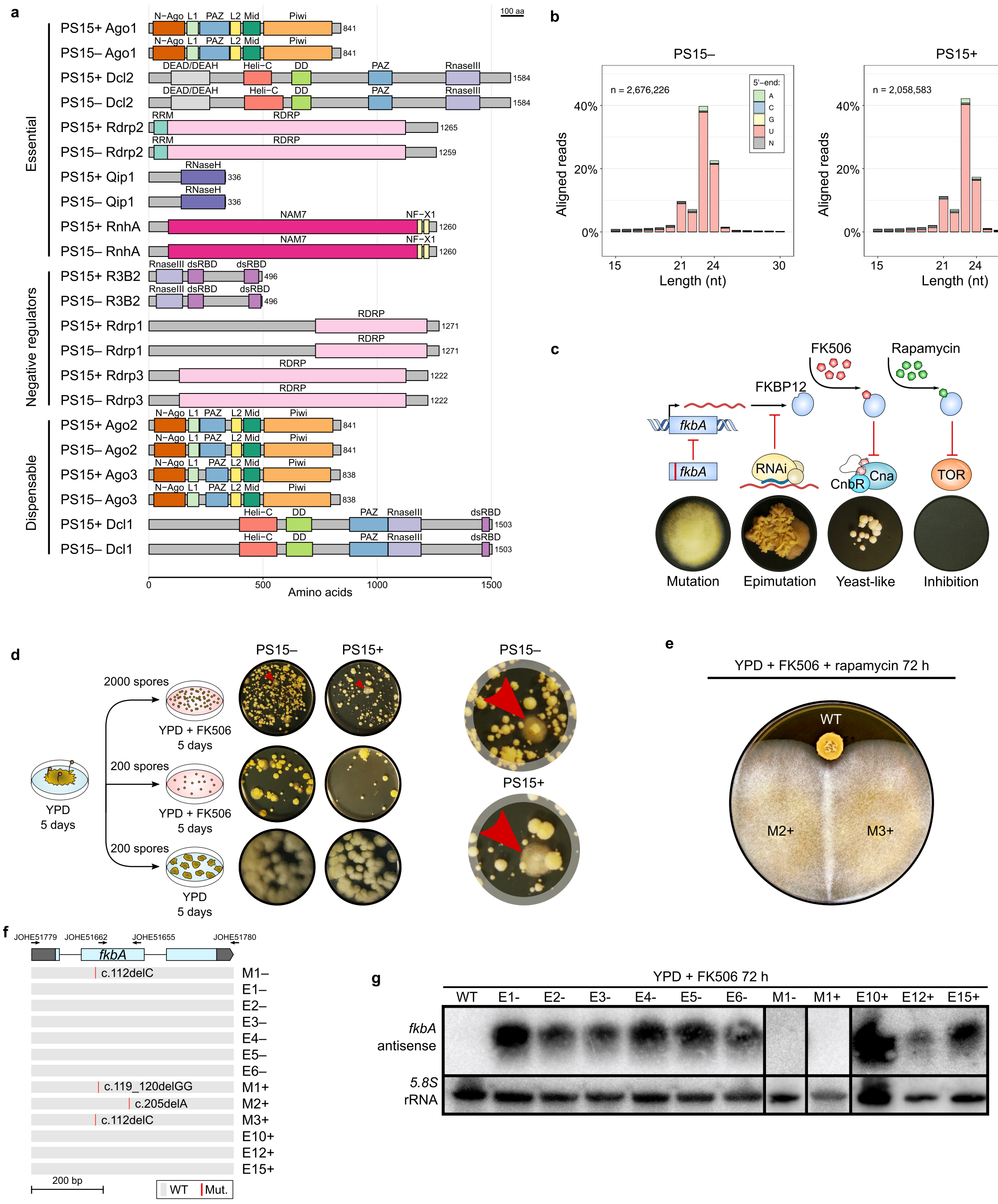


**Extended Data Fig. 1. *Mucor circinelloides* PS15 RNAi proficiency enables *fkbA* silencing. (a)** The RNAi components are classified into essential, negative regulators, and dispensable for epimutation. Rectangles display the full-length, scaled protein sequences of every identified protein homolog and their predicted, color-coded InterPro protein domains abbreviated as follows: N-Ago (Protein argonaute, N-terminal, IPR032474); L1 (Argonaute, linker 1 domain, IPR014811); PAZ (PAZ domain, IPR003100); L2 (Argonaute, linker 2 domain, IPR032472); Mid (Protein argonaute, Mid domain, IPR032473); Piwi (Piwi domain, IPR003165); DEAD/DEAH (DEAD/DEAH box helicase domain, IPR011545); Heli_C (Helicase C-terminal domain-like, IPR001650); DD (Dicer dimerization domain, IPR005034); RNaseIII (Ribonuclease III domain, IPR000999); RRM (RNA recognition motif domain, IPR000504); RDRP (RNA-dependent RNA polymerase, eukaryotic type, IPR007855); RNaseH (Ribonuclease H domain, IPR012337); NAM7 (DNA2/NAM7-like helicase, IPR045055); NF-X1 (Zinc finger, NF-X1-type, IPR000967); dsRBD (Double-stranded RNA-binding domain, IPR014720). **(b)** The percentage of sRNA aligned reads is plotted according to length and 5'-end nucleotide distribution (color-coded). **(c)** The mechanism of action of FK506 and rapamycin via FKBP12 binding and inhibition of calcineurin (Cna and CnbR) or TOR is illustrated. *fkbA* mutations (red line) or RNAi epimutations cause loss of FKBP12 function and result in drug resistance. **(d)** Isolates exhibiting loss of FKB12 function were obtained by exposing 200 or 2,000 spores to FK506 and screening for mycelium formation as an indication of resistance. Red arrows highlight representative resistant colonies, shown in the zoomed-in view on the right. **(e)** Mutant (M) isolates M2+ and M3+ exhibiting FK506 resistance were challenged with both FK506 and rapamycin for 72 hours, comparing growth to wildtype strains (WT). Isolates resistant to both drugs are labelled in black, and susceptible isolates in white. **(f)** Schematic representation of the *fkbA* gene, showing 5' and 3' untranslated regions (UTRs, dark gray blocks) and coding regions (light blue blocks). Primers used for Sanger sequencing are indicated above the gene diagram. Below, the *fkbA* nucleotide sequence is shown, with identified mutations highlighted in red as opposed to gray for wildtype sequences. **(g)** Small RNA Northern blot of depicted isolates after a 72-hour exposure to FK506, probed for antisense *fkbA* small RNAs (*fkbA* antisense, above) and sense *5.8S* ribosomal RNA (*5.8S* rRNA, below) as a loading control.

Extended Data Fig. 2

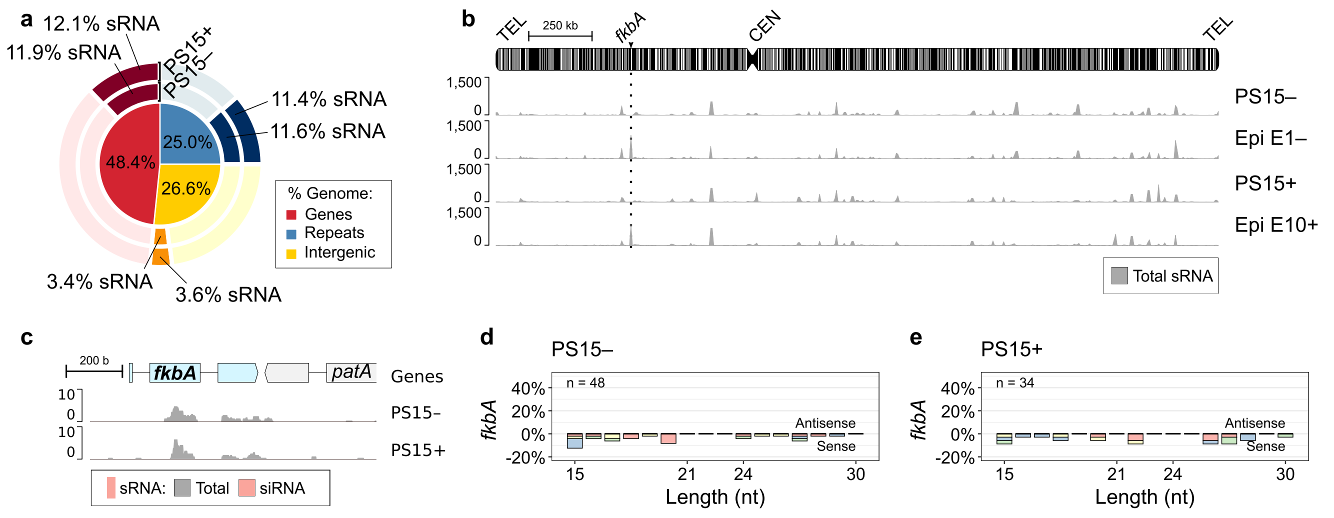


**Extended Data Fig. 2. Wildtype PS15 opposite mating types are RNAi-proficient but do not silence the *fkbA* gene in the absence of selection. (a)** Proportion of gene, repetitive, and intergenic sequences in the PS15⁻ genome (inner pie chart, color-coded). Outer pie chart shows the corresponding proportions of base pairs overlapped by sRNA loci (darker shade) and not overlapped (lighter shade). **(b)** Genome-wide distribution of total sRNA (dark gray) across the chromosome carrying *fkbA* (arrowhead and dotted line) in wildtype, naive PS15– and PS15+, as well as representative epimutants of each mating type. Repeats are shown as black vertical lines; the centromere (CEN) as a constriction; telomeres (TEL) as rounded ends. **(c)** sRNA coverage (dark gray) and siRNA (red) across the *fkbA* locus in PS15– and PS15+. **(d, e)** Length distributionand 5'-end nucleotide composition (color-coded) according to strand sense of sRNA reads mapping to *fkbA* in wildtype, naive PS15– (d) and PS15+ (e).

Extended Data Fig. 3

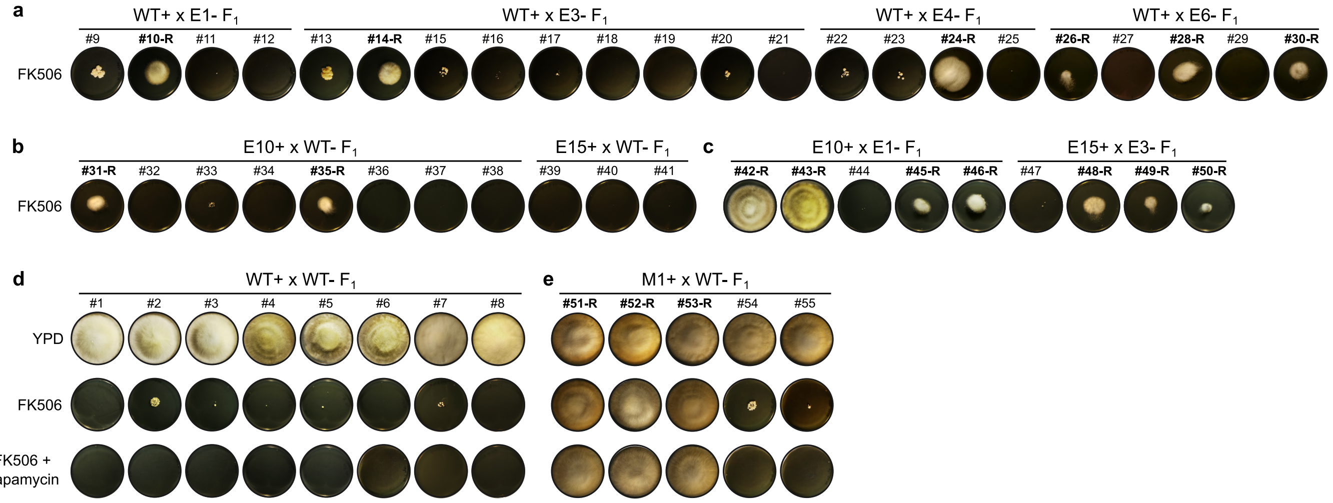


**Extended Data Fig. 3. Independent FK506 challenge and $F_1$ progeny derived from control crosses. (a-c)** FK506 susceptibility was assessed in the progeny from **(a)** – mating type epimutant crossed with + mating type wildtype strain; **(b)** + mating type epimutant crossed with – mating type wildtype strain; and **(c)** a cross between two epimutants (– and +). This independent challenge with FK506 revealed the same susceptibility patterns as shown in Fig. 2a-c for FK506 and rapamycin combined exposure, indicating the observed resistance was already present in the germspores before the drug challenge, as opposed to arising de novo due to treatment. **(d, e)** Susceptibility to FK506 alone and in combination with rapamycin was determined in the progeny from **(d)** a cross between two wild type (– and +); and **(e)** – mating type *fkbA* mutant crossed with + mating type wildtype strain. Growth in YPD served as a viability control.

Extended Data Fig. 4

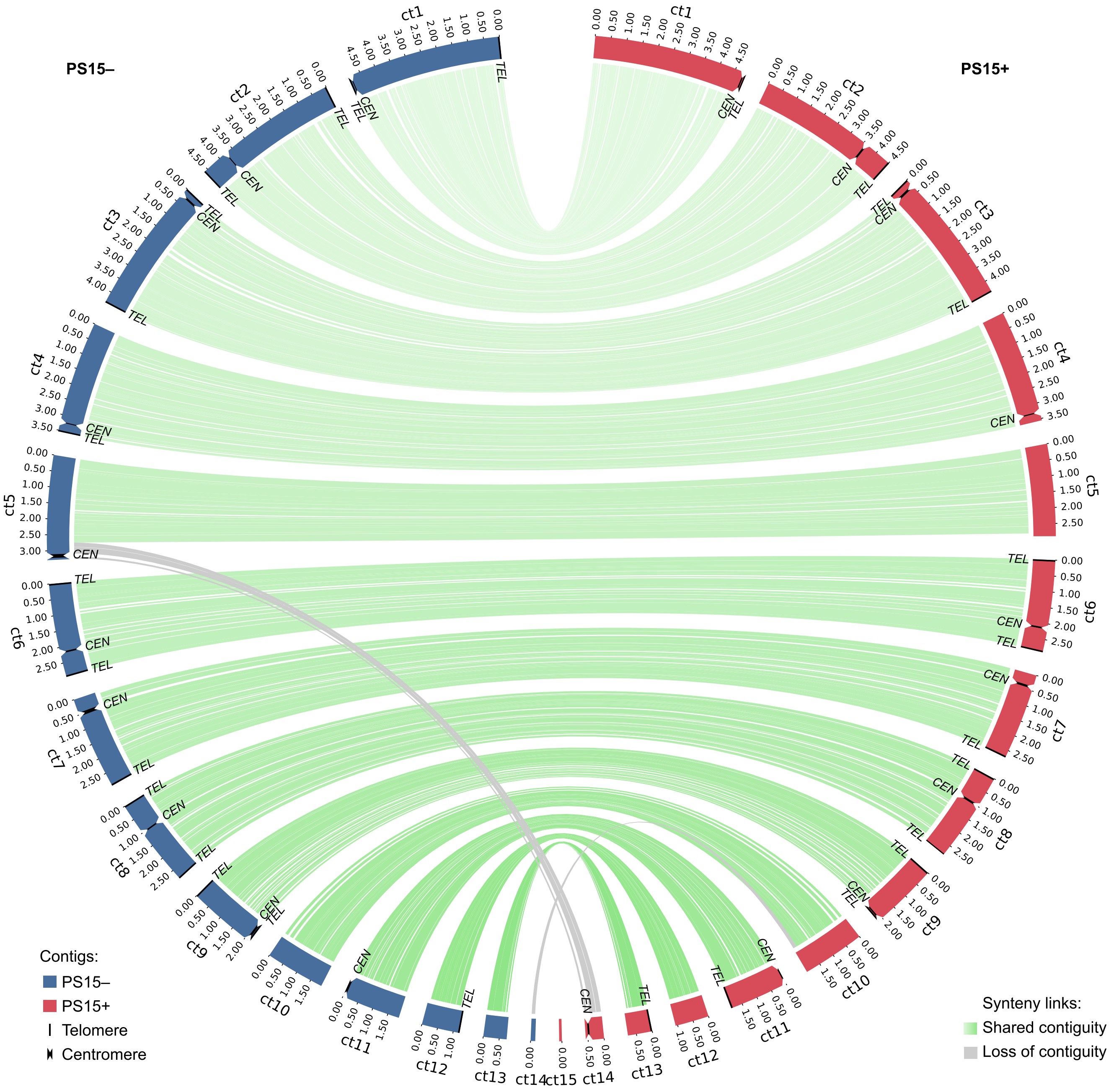


**Extended Data Fig. 4. *Mucor circinelloides* PS15 – and + genome assemblies.** Contig distribution and length of each assembly are shown as colored rectangles (– in blue, + in red). Centromeres (*CEN*) are depicted as constrictions and telomeres (*TEL*) are indicated by black lines closing the contig. Synteny between homologous contigs from each strain is shown as green lines. In total, 13 shared synteny blocks were defined and used in meiotic recombination and coverage analyses shown in Extended Data Fig. 5 and 6, comprising six complete telomere-to-telomere chromosomes (ct1, ct2, ct3, ct6, ct8, and ct9) and seven contigs (ct4, ct5, ct7, ct10, ct11, ct12, ct13), as well as nine centromeres located on ct1, ct2, ct3, ct4, ct6, ct7, ct8, ct9, and ct11. On the other hand, gray lines indicate similarity between regions that are not contiguous in both strains and were excluded from further meiotic recombination and coverage analyses. One centromere lies outside the defined synteny blocks. In PS15–, it is located on ct5, whereas in PS15+, the homologous region is split between ct14 (which harbors the centromere) and ct5. Due to unresolved chromosomal linkage between ct14 and ct5 in PS15+, the centromere-containing region was excluded from the synteny blocks used in recombination analyses.

### Extended Data Fig. 5

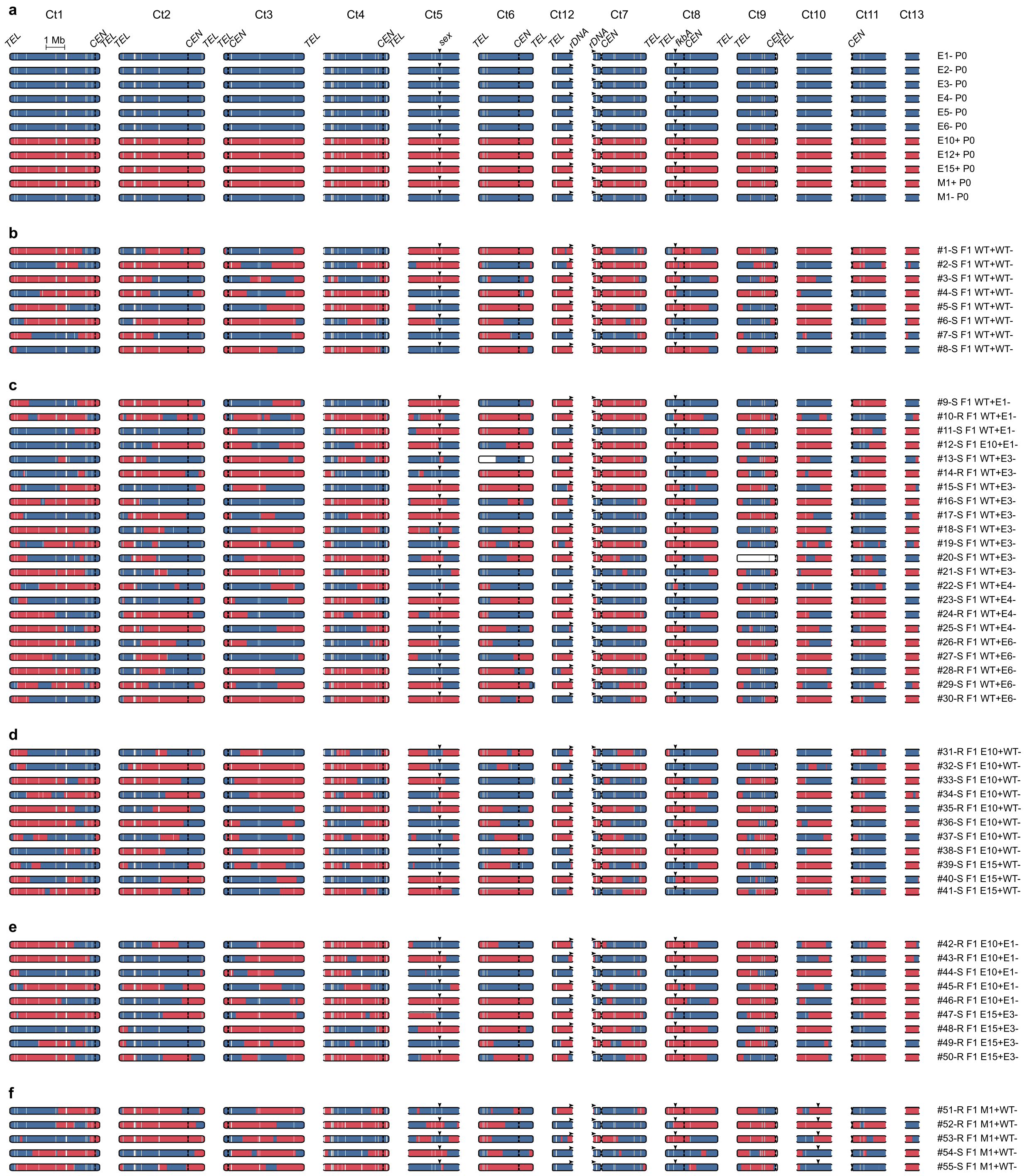


**Extended Data Fig. 5. Meiotic recombination sites in $F_1$ progeny. (a-f)** Contigs exhibiting synteny and contiguity in both PS15– and + are shown to indicate meiotic recombination sites in: **(a)** parental strains ($P_0$) utilized for sexual crosses, and $F_1$ progeny derived from **(b)** a cross between two wild type (– and +); **(c)** – mating type epimutant crossed with + mating type wildtype strain; **(d)** + mating type epimutant crossed with – mating type wildtype strain; **(e)** a cross between two epimutants (– and +); and **(f)** – mating type *fkbA* mutant crossed with + mating type wildtype strain. The contig (Ct) drawings depict the inheritance pattern of genetic variation identified in both opposite mating types (SNPs, – mating type in blue and + mating type in red). Incomplete chromosomes, i.e., without telomeric repeats are depicted as rounded rectangles with open ends, whereas full chromosomes show a centromere (constriction) and telomeric repeats on both ends (closed ends). Relevant loci are designated by arrows (RFLPs, rDNA, *sex*, and *fkbA*), indicating their direction (forward or reverse) when relevant (rDNA). To facilitate visualization, Ct12 is shown horizontally inverted and placed to the left of Ct7 to suggest that they are most likely joined at the rDNA repeats, as both contig ends show genetic linkage in all progeny.

Extended Data Fig. 6

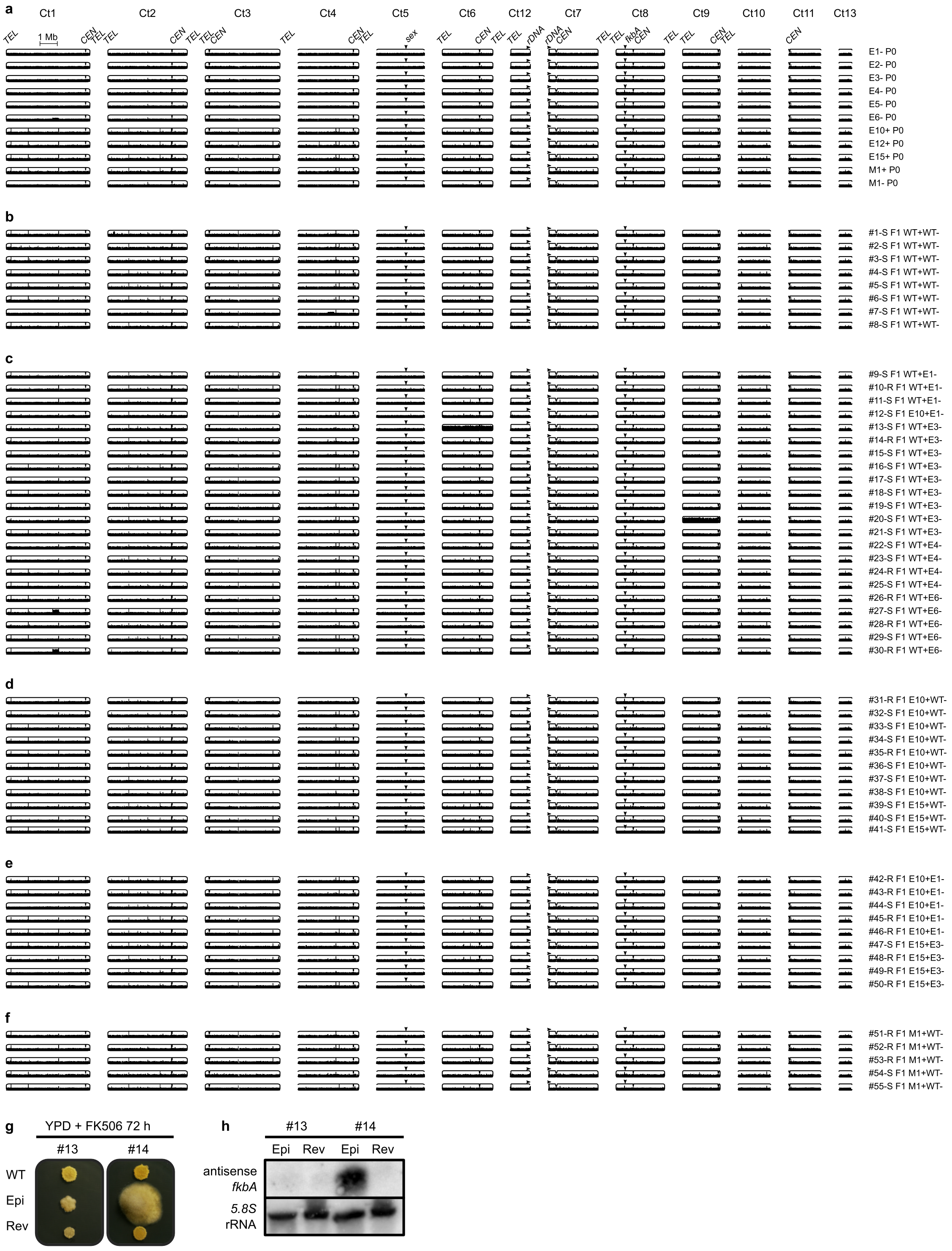


**Extended Data Fig. 6. Changes in ploidy or segmental duplications or deletions are not linked to RNAi epimutations. (a-f)** Isolates, contigs, and their features are arranged as indicated in Extended Data Fig. 5. To facilitate visualization of changes in ploidy, DNA coverage was normalized as counts per million. An E6- subpopulation harbored a segmental duplication in contig 1 (Ct1), which was inherited by two progeny (isolates #27 and #30). This duplication was unlinked to changes in FK506 susceptibility. In addition, progeny isolates #13 and #20 showed chromosome 6 (Ct6) and 9 (Ct9) duplication, respectively. Ct9 duplication does not result in FK506 and/or rapamycin resistance. **(g)** Progeny #13 (tolerant, Ct6 duplication) and #14 (resistant, euploid) were passaged on non-selective medium for ten 72-hour passages (revertants) and transferred onto FK506-selective medium next to their corresponding wildtype susceptible control and resistant isolates to assess reversion. **(h)** Small RNA Northern blot of progeny isolates #13 and #14 after a 72-hour exposition to FK506. The blot shows probe hybridizations to detect antisense *fkbA* small RNAs (*fkbA* antisense, above) and sense *5.8S* ribosomal RNA (*5.8S* rRNA, below) as a loading control. Isolate #13 did not harbor sRNAs. Although chromosome 6 duplication may confer tolerance to FK506 and rapamycin, tolerant isolate #13 exhibits yeast-like morphology and lacks antisense sRNAs targeting *fkbA*.

Extended Data Fig. 7

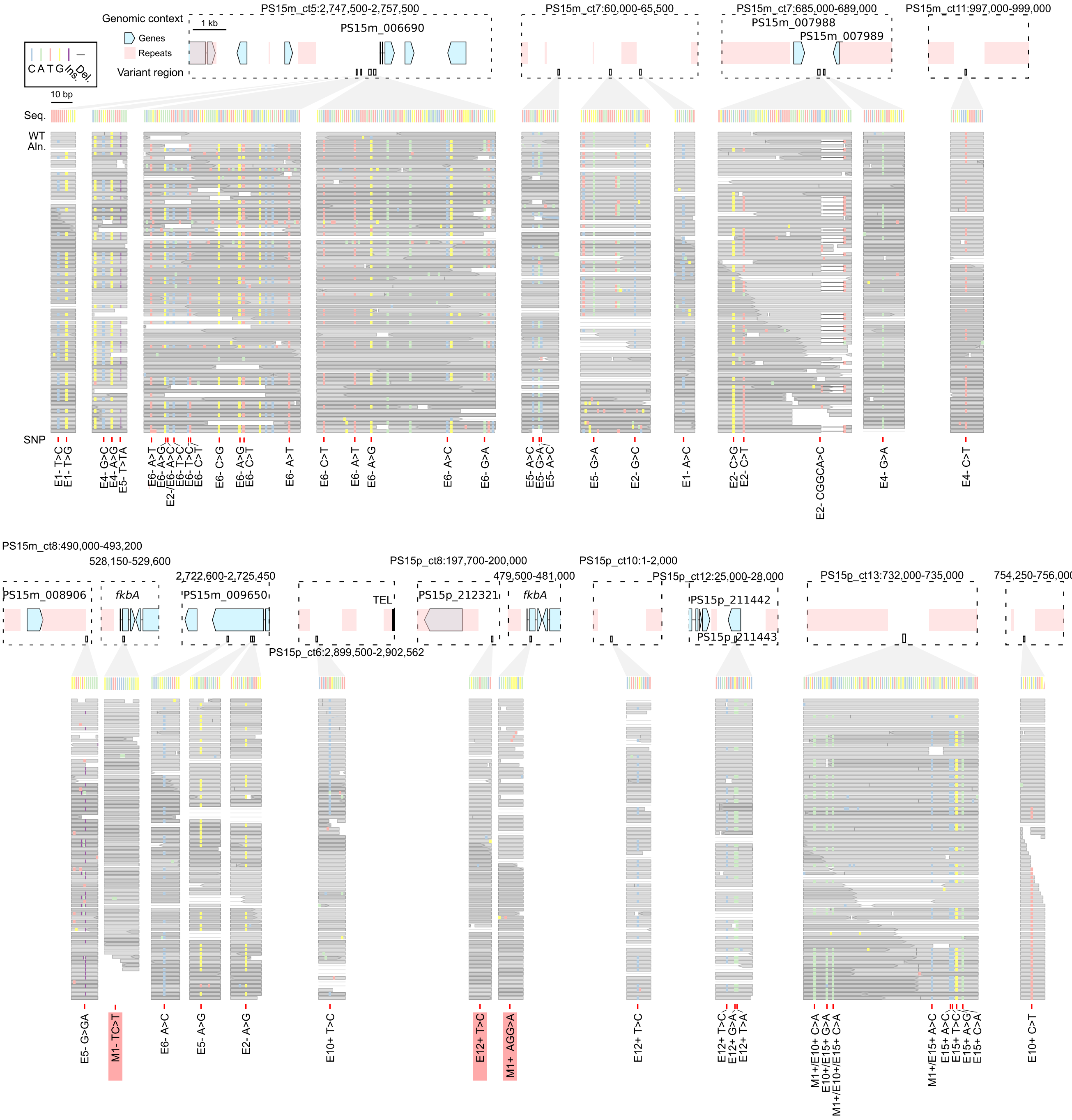


**Extended Data Fig. 7. Whole-genome sequencing reveals no consistent mutations conferring FK506 and rapamycin resistance beyond the RNA-silenced, epimutant *fkbA* locus.** Putative single nucleotide polymorphisms (SNPs, red) are shown within their genomic context (top; coordinates indicated). Genes are represented as light blue arrowed blocks, repeats as light red blocks, and a telomeric region (TEL) as a black block. Genes nearest to each variant are labeled. Below, zoomed-in panels (corresponding to boxed regions above) display the reference sequence (Seq.) and wildtype read alignments (WT Aln.), both color-coded as indicated. Wildtype read alignments show pre-existing variants to assess whether putative SNPs are false positives or bona fide mutations. Variants not reproduced in the wildtype alignment reads, regarded as bona fide mutations, are highlighted with red shading.

Extended Data Fig. 8

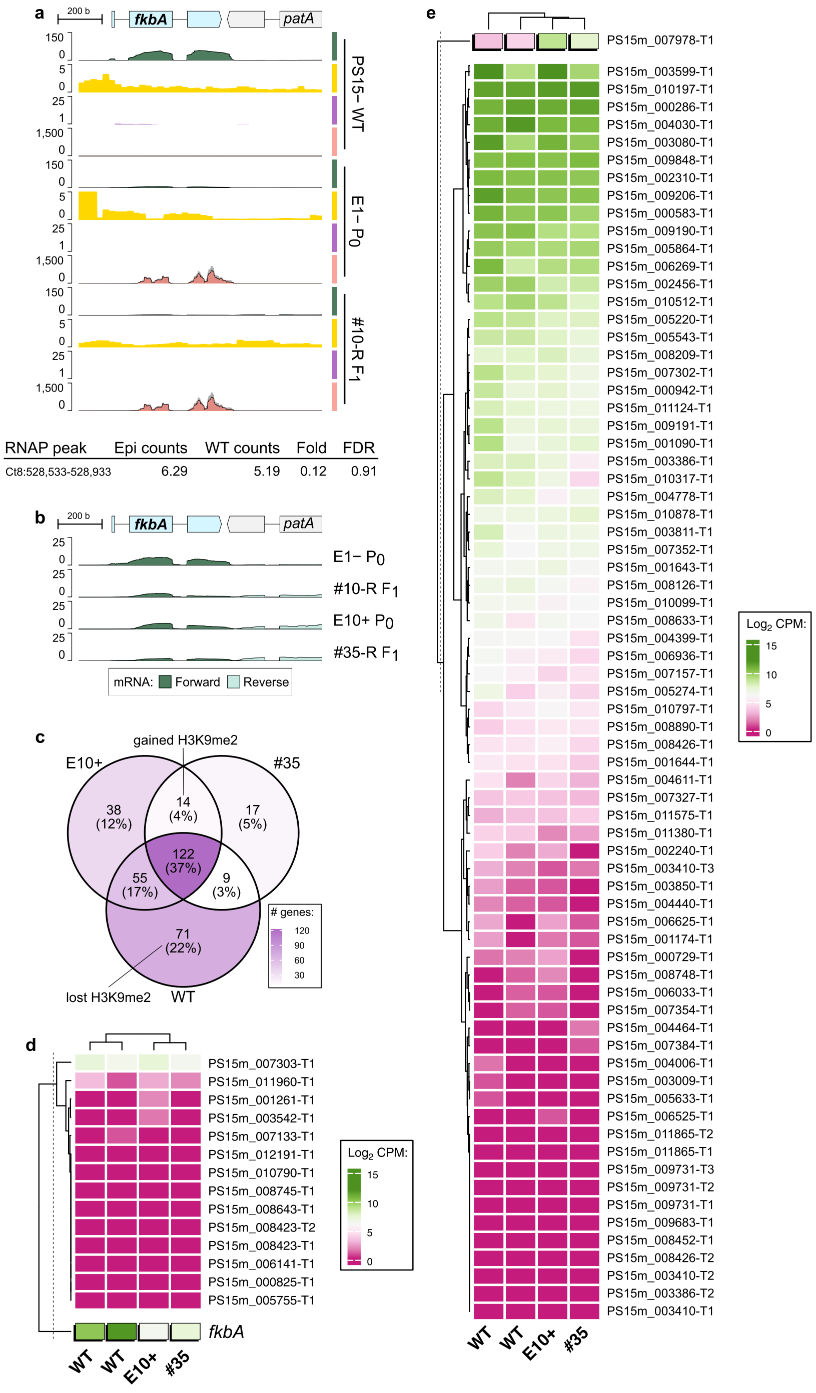


**Extended Data Fig. 8. Epimutations are consistent with posttranscriptional gene silencing.** **(a)** Genomic tracks showing H3K9me2 enrichment (purple), RNA Polymerase II enrichment (RNAP, yellow), stranded mRNA (green; forward and reverse overlaid), and sRNA (dark gray for total sRNA, overlaid with red for siRNA) in *M. circinelloides* PS15– wildtype, the epimutant parent (E1– P0), and its F1 progeny (#10-R F1) across an expanded view of the *fkbA* locus. Below, differential binding analysis (DiffBind, DESeq2) reveals no significant differences in RNAP enrichment between epimutant and wildtype isolates (Fold enrichment = 0.12, FDR = 0.91). **(b)** Stranded mRNA across the *fkbA* locus displayed at a lower CPM range to highlight subtle *fkbA* expression in epimutants and their progeny from both mating types. **(c)** A Venn diagram shows overlapping genes embedded in H3K9me2 ($\geq 0.05$ 90% of their coding sequence) in a naïve wild-type strain (WT), the epimutant E10+, and its progeny (#34). Color-coded shades of purple indicate the number of overlapping genes. Gained and lost H3K9me2 genes are highlighted, defined as those embedded in both epimutant and progeny but absent in WT (gained), or present in WT only (lost). **(b, c)** The heatmaps display color-coded gene expression values as normalized $\log_2$ CPM across two wildtype duplicates, an epimutant (E10+), and its progeny (#34), which served as duplicates in the differential expression analysis (DeSEQ2). Significant expression changes ($\text{FDR} \leq 0.05$) between wild type and epimutant are indicated by raised, black-shadowed cells. The similarity between gene expression values and samples is depicted by rows and columns dendrograms generated through complete clustering using the Euclidean distance matrix. A dashed vertical line indicates a split in the dendrogram node, which was used to mark the positive control gene. In **(d)**, fourteen H3K9me2 gained genes are displayed, using *fkbA* as a positive control for expected expression values due to gene silencing. In **(e)**, 71 H3K9me2 lost genes are shown, using *PS15m_007978* as a positive control of an upregulated gene. Except for the positive controls, none of these genes showed significant expression changes as a consequence of gaining or losing H3K9me2.

Extended Data Fig. 9

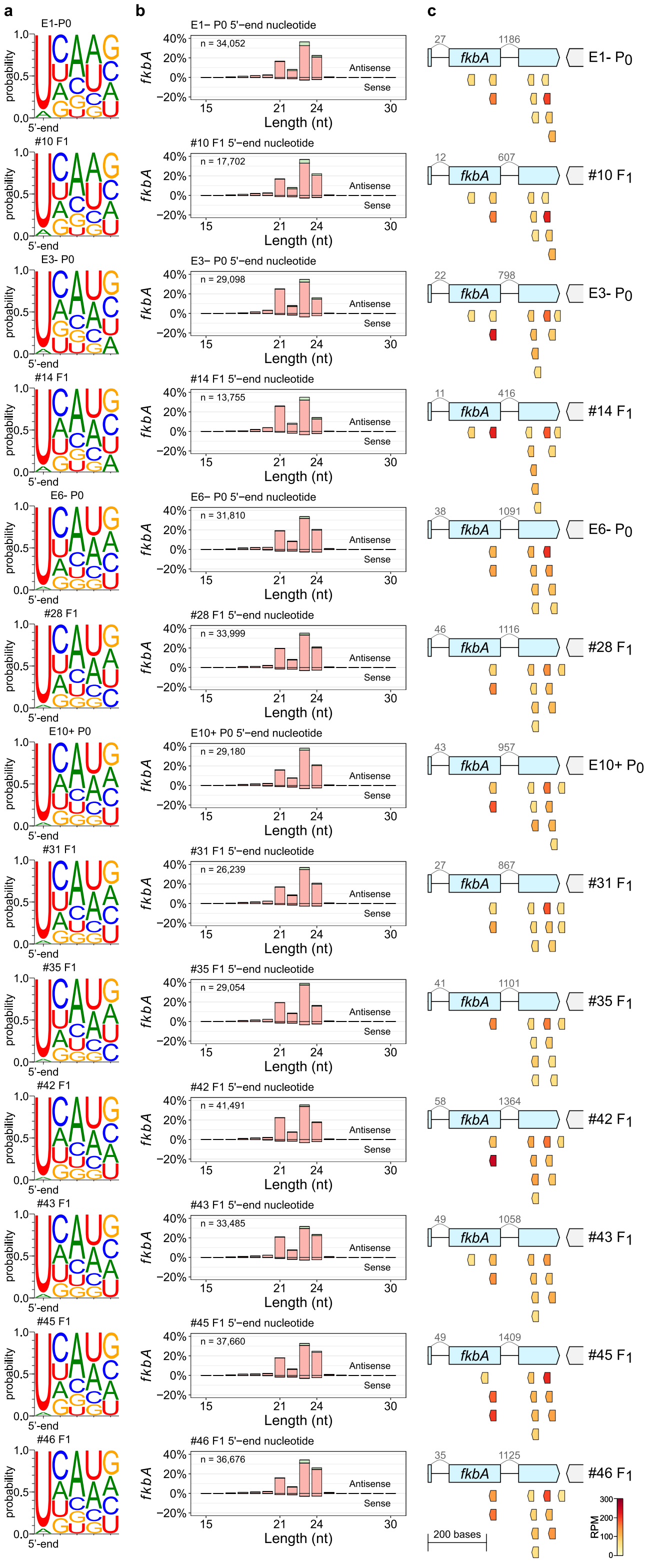


**Extended Data Fig. 9. Inheritable siRNAs harbor common features and the most abundant molecules can be traced from progeny to their parents. (a)** Sequence logos display the proportion of each nucleotide (color-coded) at the first five (5'-end) positions of the siRNA sequences found in the depicted epimutant parents and progeny. **(b)** The percentage of siRNA reads is plotted according to length, strand sense, and 5'-end nucleotide distribution. **(c)** The ten most abundant siRNA molecules in each epimutant are color-coded according to their relative abundance as reads per million (RPM) and plotted across the *fkbA* locus. In the gene model, the number of siRNA reads spanning across each exon-exon junction are displayed above curved lines (in grey) for each epimutant. Inheritable siRNA molecules exhibit lengths ranging from 21 to 24 nt, harbor a 5'-uracil, are antisense, and derive mainly from mature mRNAs. There is a notable correlation between the most abundant siRNAs found in parents and their progeny.

#### Extended Data Fig. 10

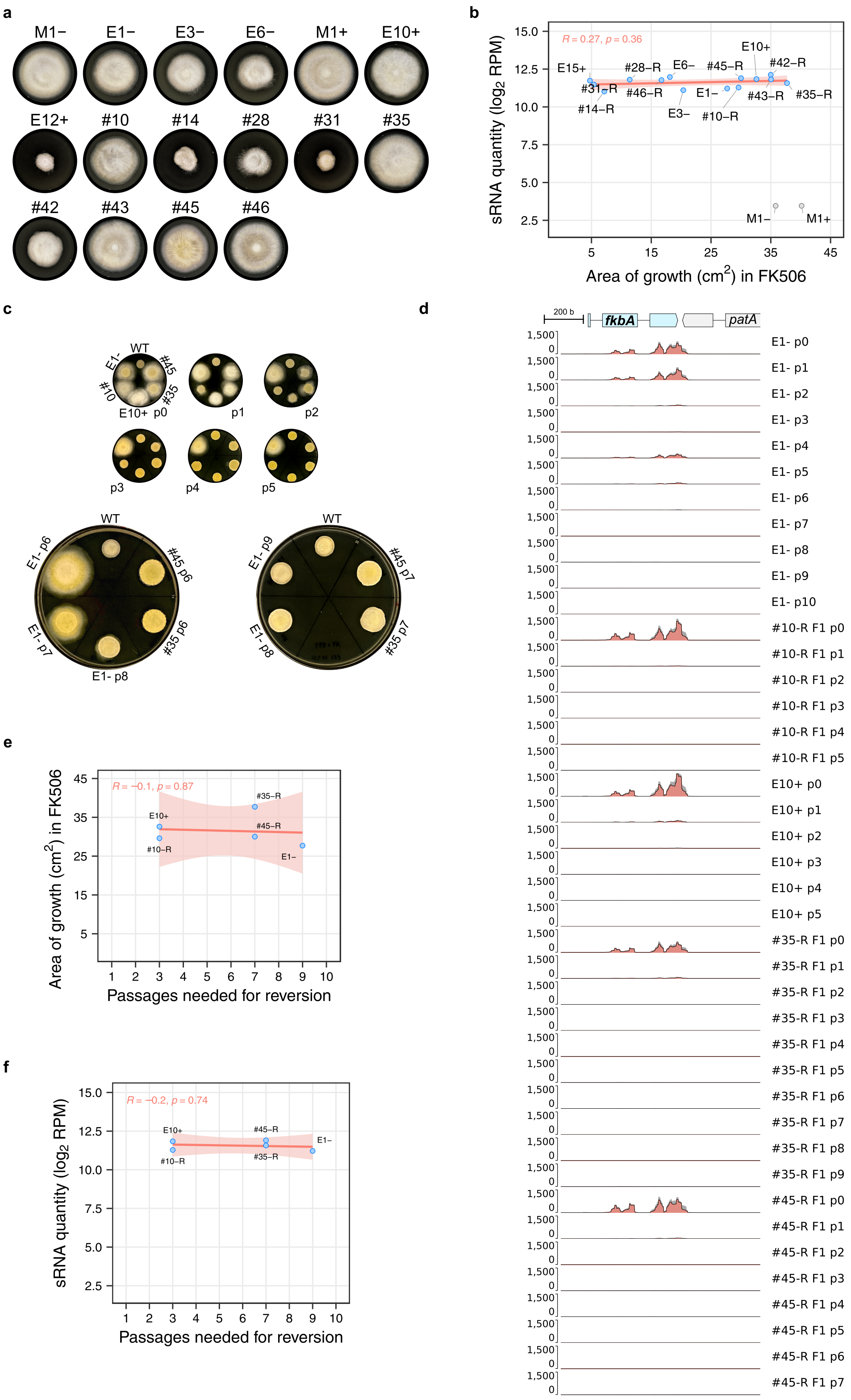

**Extended Data Fig. 10. Progressive loss of FK506 and rapamycin resistance coincides with siRNA depletion. (a)** Growth of representative resistant strains during 72-hour exposure to FK506 and rapamycin. **(b)** Dot plot showing the correlation between growth area (cm²), used as a proxy for the extent of resistance, and sRNA quantity, measured as normalized $log_2$ reads per million (RPM). A red regression line with a shaded confidence interval is shown. Pearson's R and the associated p-value are indicated in red. Although resistance levels vary among isolates, they do not correlate with sRNA quantity. **(c)** Growth of selected strains during FK506 exposure across successive drug-free vegetative passages until reversion to drug susceptibility. Passage number increases from left to right. **(d)** Genomic plot showing sRNA coverage (dark gray for total sRNA, red overlay for siRNA) across the *fkbA* locus in the corresponding drug-free passages shown in **(c)**. **(e, f)** Dot plots showing correlations between the number of passages required for reversion and the extent of resistance (**e**) or sRNA quantity (**f**) in strains shown in **(d)**. A red regression line with a shaded confidence interval is shown. Pearson's R and the p-value are indicated in red.
