## Supplementary figures and images for "RNAi epimutations conferring antifungal drug resistance are inheritable"

### sourceDataExtDataFig1gRibosomalRNANorthernBlot.tif

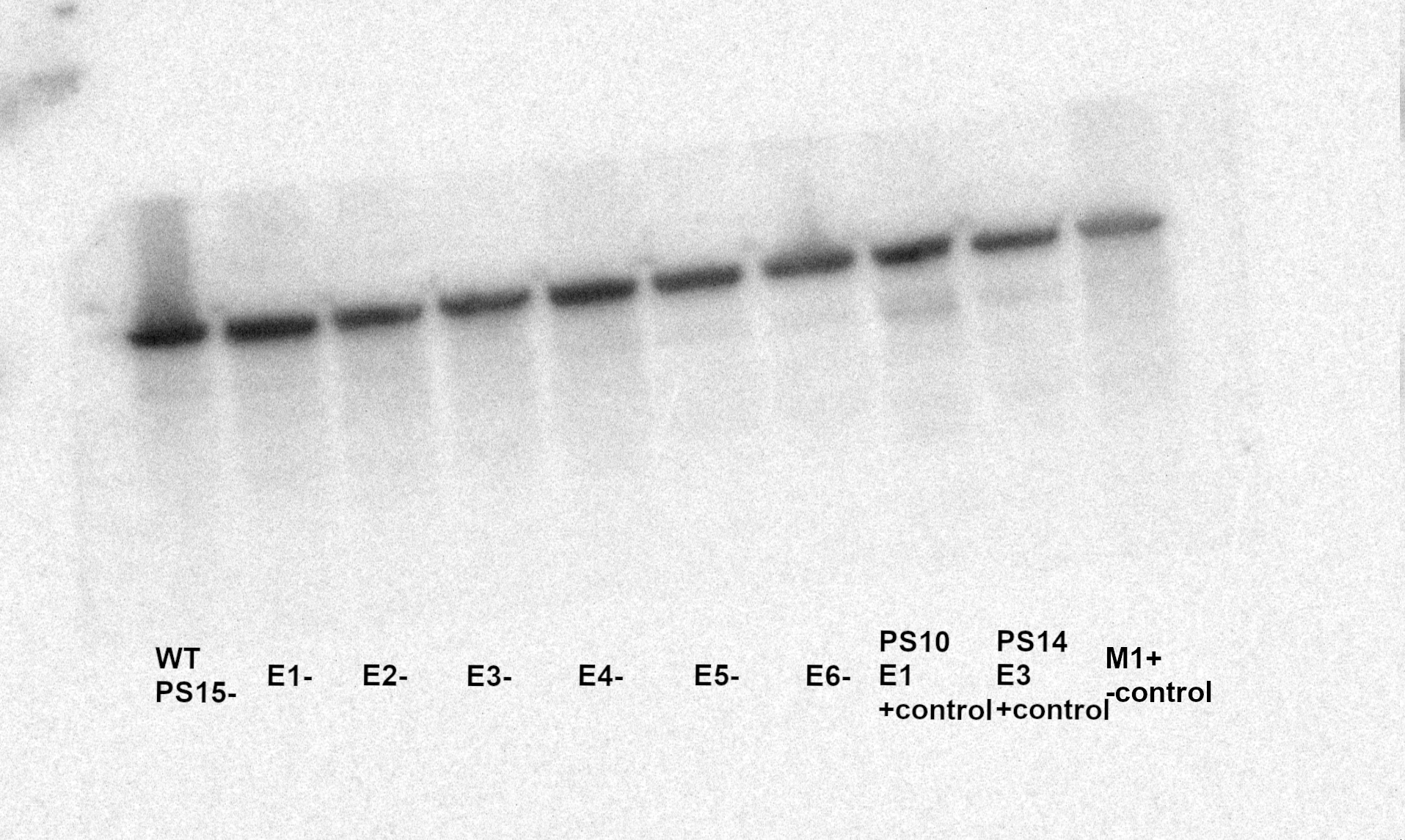
